# The NDH complex reveals a trade-off that constrains maximising photosynthesis in *Arabidopsis thaliana*

**DOI:** 10.1101/2022.11.13.516254

**Authors:** Tom P.J.M. Theeuwen, Aaron W. Lawson, Dillian Tijink, Federico Fornaguera, Frank F.M. Becker, Ludovico Caracciolo, Nicholas Fisher, David M. Kramer, Erik Wijnker, Jeremy Harbinson, Mark G.M. Aarts

## Abstract

The Green Revolution has resulted in major improvements in crop productivity, but left photosynthesis largely unimproved. Despite ample variation of photosynthetic performance in crops and their wild relatives, the photosynthetic capacity of elite breeding lines remains well below its theoretical maximum. As yield is often the primary selective trait, current plant breeding approaches result in photosynthetic trade-offs that prevent positive selection for photosynthetic performance itself. Currently, genetic variation for photosynthetic performance is seldomly validated at the genetic level, and as a result these photosynthetic trade-offs remain poorly understood. Here we reveal the physiological nature of a photosynthetic trade-off caused by the NAD(P)H dehydrogenase (NDH) complex. The use of an *Arabidopsis thaliana* cybrid panel revealed how a natural allele of the chloroplastic gene *NAD(P)H-QUINONE OXIDOREDUCTASE SUBUNIT 6 -* a subunit of the NDH complex - results in a faster recovery of photosystem II efficiency after a transition from high to low irradiances. This improvement is due to a reduction in NDH activity. Under low-light conditions this reduction in NDH activity has a neutral effect on biomass, while under highly fluctuating light conditions, including high irradiances, more NDH activity is favoured. This shows that while allelic variation in one gene can have beneficial effects on one aspect of photosynthesis, it can, depending on environmental conditions, have negative effects on other aspects of photosynthesis. As environmental conditions are hardly ever stable in agricultural systems, understanding photosynthetic trade-offs allows us to explore shifting photosynthetic performance closer to the theoretical maximum.

## Introduction

The Green Revolution resulted in unprecedented increases in crop yields, however, yield increases have stagnated in recent years in the face of looming global food insecurity (Zhu et al., 2010; Ray et al., 2012). Plant breeding contributed to these yield increases by focusing on improving the light interception index and the harvest index, but has left the efficiency of converting absorbed light energy into biomass, primarily determined by photosynthesis, largely unimproved (Monteith, 1994; Long et al., 2006). In fact, for most staple crops the increase in photosynthetic efficiency has been minimal since the Green Revolution (Long et al., 2015; Theeuwen et al., 2022b). This poses a conundrum; while it has been shown that there is evidence of ample variation for photosynthetic performance in elite breeding lines, this variation does not correlate well with higher yields, the trait often selected for in plant breeding (Driever et al., 2014; Acevedo-Siaca et al., 2020). In part, this discrepancy could be explained by the sheer amount of Quantitative Trait Loci (QTLs) responsible for the variation in photosynthetic performance (Van Rooijen et al., 2017; Oakley et al., 2018; Prinzenberg et al., 2020), where alleles of QTLs can even have opposing or interacting effects, making selection on QTLs for photosynthesis via yield complicated. However, another important factor is that variation with a positive effect on photosynthesis under some conditions might have, under other conditions, negative effects on photosynthesis, or have negative effects on yield due to other consequences of improved photosynthesis. Examples of photosynthetic trade-offs include how photo- protective mechanisms are essential, but come at a cost of light use efficiency (Rutherford et al., 2012), and how for Rubisco increasing specificity for CO_2_ relative to O_2_ limits the catalytic rate and vice versa (Zhu et al., 2004; Noor and Milo, 2012; Flamholz et al., 2019). Revealing and understanding more of such photosynthetic trade-offs is essential for pushing photosynthetic performance closer to the theoretical maximum and in turn improving yields.

Modelling photosynthesis has revealed a number of targets for improved photosynthetic performance. One of these targets is the aforementioned trade-off between photo-protective mechanisms and light use efficiency. However, photo-protection mechanisms adapt relatively slow to fluctuating light conditions, which can reduce the potential assimilation by up to 30% (Long et al., 2015, 2022; Slattery et al., 2018; Tanaka et al., 2019). Light conditions are rarely stable in field crop systems; passing clouds produce so called cloud flecks, while leaf movements inside canopies produce so called sun flecks (Kaiser et al., 2018; Durand et al., 2021). These fluctuations in irradiance result in specific problems for photosynthesis. Rapid increases in irradiance frequently result in a greater problem of excess light, because the increase in photosynthesis that would normally occur in response to an increase in irradiance is relatively slow. The engagement of photo-protective mechanisms in response to excess light can act to thermally dissipate the excess of excited states of chlorophyll formed in PSII and PSI resulting in less photodamage. These photo- protective mechanisms are referred to as non-photochemical quenching (NPQ). NPQ is triggered in response to a decrease in the pH of the thylakoid lumen, a change which occurs once the capacity of photosynthetic electron transport exceeds the activity of the photosynthetic metabolism. This fall in lumen pH results in an enhanced difference in the transthylakoid pH, which together with the transthylakoid voltage, is also a driving force for ATP synthesis and thus for photosynthetic metabolism. The decrease in lumen pH results in protonation of PsbS and activation of the xanthophyll cycle, which together induce, or induce further if it is already active, the fast-relaxing component of NPQ known as *q*_E_ that protects PSII. The fall in lumen pH also slows electron transport through the cytochrome *b_6_f* complex. This serves to protect PSI by limiting electron transport on the donor side of PSI, reducing the risk of acceptor side limitation which brings with it the risk of photodamage to PSI. However, a subsequent reduction in light results in a relatively slow relaxation and excess activity of *q*_E_, in turn leading to an overly down-regulated PSII, which limits assimilation. Using genetic modification approaches to overexpress *PsbS* and the enzymes involved in the xanthophyll cycle in tobacco indeed resulted in faster relaxation of NPQ and significant increases in biomass (Kromdijk et al., 2016). The same approach in soybean likewise resulted in increased biomass, but only did so in one of the two growing seasons tested (De Souza et al., 2022). In *Arabidopsis thaliana* this approach did not result in the faster relaxation of NPQ, nor to an increase in biomass (Garcia-Molina and Leister, 2020), indicating it is not a ‘one-size fits all’ solution.

In nature, dynamic light conditions are everywhere, but their frequency, the difference between subsequent light levels and the combination with other environmental conditions might favour photo- protection characteristics that are environment specific. This also means trade-offs preventing certain characteristics in one environment might be absent or completely different in another environment. This may be the reason there is ample natural variation for the capacity to deal with dynamic light conditions (Rungrat et al., 2019; Tanaka et al., 2019; Acevedo-Siaca et al., 2021) and other aspects of the operation with regulation and optimisation of photosynthesis not directly connected with fluctuating light. In the case of dynamic properties of NPQ unveiling the genetic variation, i.e. genes and alleles, outlining these differences will generate insights into the physiological processes underlying NPQ and open up novel targets for improving photosynthesis (Theeuwen et al., 2022b). Up to 3000 proteins are involved with photosynthetic functioning, most of which are nuclear encoded and targeted to the chloroplast. All this while the organellar genomes encode only roughly 150 proteins (Timmis et al., 2004), those of which the majority are for its own transcription and translation machinery, leaving the chloroplast to encode roughly 40 proteins involved in photosynthesis and the mitochondria roughly 14 proteins involved in the oxidative chain (Westhoff et al., 1998). However, while the organelles only encode a fraction of the proteins involved in photosynthesis and related processes, those proteins form a central hub in the overall photosynthetic machinery (Joseph et al., 2013; Budar and Roux, 2014). Studying variation in the organellar genomes, collectively referred to as the plasmotype, is difficult due to the inability of separating their effect from nuclear variation, termed the nucleotype (Budar and Roux, 2011). Systematically screening natural variation for the dynamic properties of NPQ, encoded by the organellar genomes, is not trivial, and only recently has an efficient method been developed using a maternal haploid induction system to generate novel cybrids of *A. thaliana* on a large scale (Flood & Theeuwen et al., 2020).

Here we report the high-throughput screening of a new cybrid panel, which contains species-wide organellar diversity in *A. thaliana* for photosynthetic responses to fluctuating light. Grown in semi-protected conditions for two consecutive years, the cybrid panel revealed a specific plasmotype that showed lower NPQ after a transition to low-light, and this coincided with a capacity for a quicker recovery of photosynthesis after fluctuations in light conditions. Using a genetic exclusion strategy, we pin-pointed a single substitution of a highly conserved amino acid in NAD(P)H-QUINONE OXIDOREDUCTASE SUBUNIT 6 (NDHG) as the causal variant. The phenotypic difference between the alleles revealed a direct link between NDH and recovery of the light-use efficiency for electron transport by photosystem II after transition from high-light to low-light conditions. Lastly, this allelic variant reveals the NDH complex to be part of a photosynthetic trade-off where a potentially positive effect on photosystem II does not translate to higher shoot dry weight due to limitations imposed by the reduced cyclic electron transport, those of which are especially pronounced in highly fluctuating light conditions.

## Results

### Natural variation for dynamic photosynthesis in organellar genomes

As the distribution of *A. thaliana* is spread over large sections of the Eurasian land mass and high altitude regions of Africa (Alonso-Blanco et al., 2016; Durvasula et al., 2017; Zeng et al., 2017; Zou et al., 2017) different accessions have been exposed to a diverse range of environments. It is, therefore, likely that in such diverse environmental conditions, different adaptations to dynamic environments have occurred. The combination of 60 diverse, but globally representative plasmotypes, with four diverse nucleotypes (Theeuwen et al., 2022a), resulted in a novel cybrid panel that we used to explore the dependency of variation of dynamic photosynthetic properties on variation of the plasmotype. The cybrid panel was grown in semi-protected cultivation conditions in two consecutive years during the springs of 2020 and 2021 in Wageningen, the Netherlands (51°59’20.0“N 5°39’43.2”E). The plants were grown in an unheated, gauze covered greenhouse in deep pots (Figure 1A), allowing them to be moved for high-throughput phenotyping. Both years were characterized by a relatively cold first week of the experiment. In 2020 the latter part of the growing season saw predominantly sunny days while in 2021 the weather was half cloudy with intermittent sun (see Supplementary Figure 1 and 2 for temperature and light irradiances throughout the experiment).

**Figure 1.**
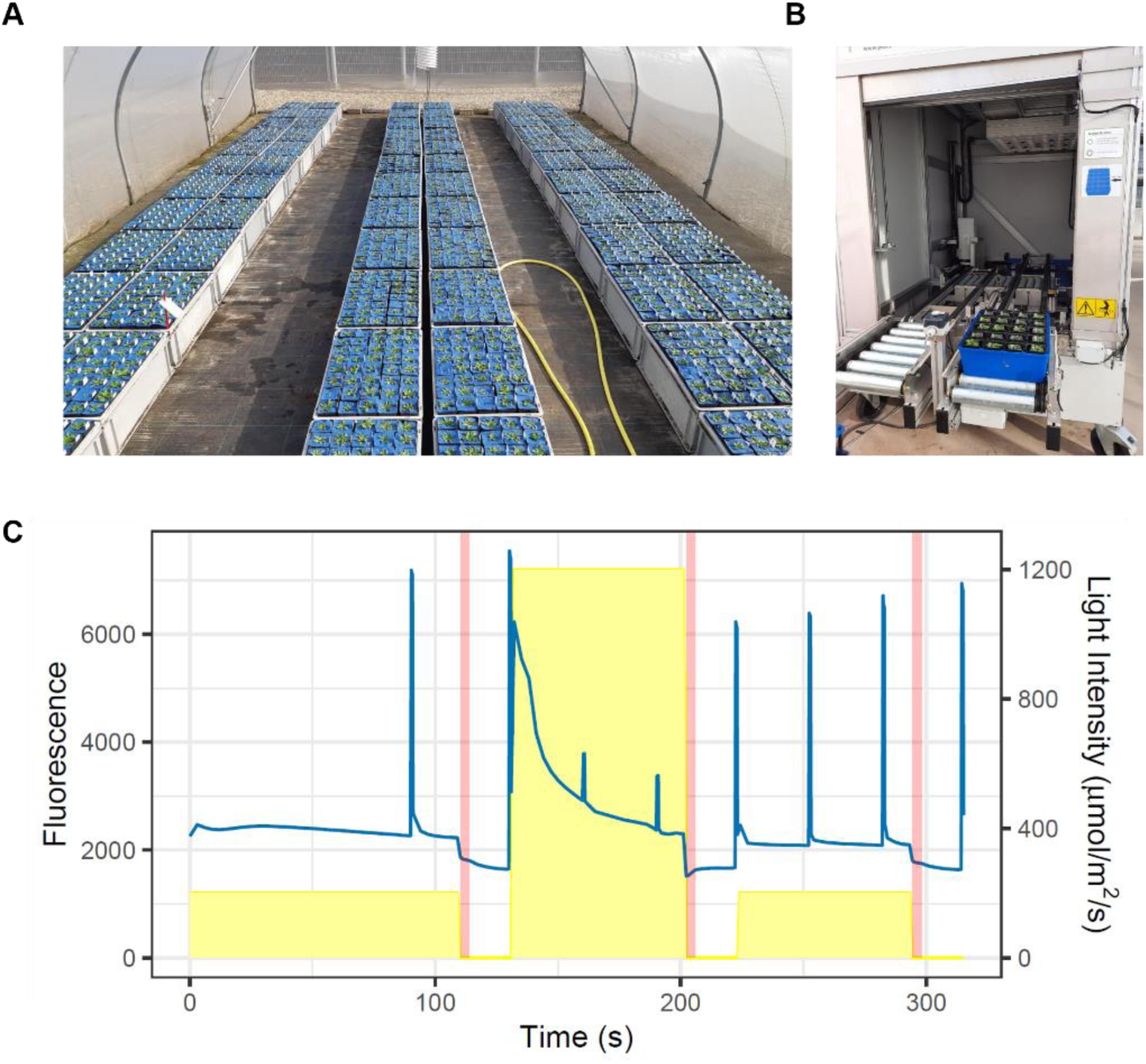
Overview of cybrids grown in semi-protected cultivation conditions and the high-throughput phenotyping method. A) Impression of the gauze tunnel experiment in Spring 2022. B) Impression of the high-throughput chlorophyll fluorescence imaging system. C) Six minute light regime (yellow) and chlorophyll fluorescence measuring regime was used to screen the cybrid panel. A two second far-red period is given (red) after all three light periods. The chlorophyll fluorescence signal (blue) of one representative plant is shown to depict when the saturating pulses were applied to calculate the photosynthetic parameters.

**Figure 2.**
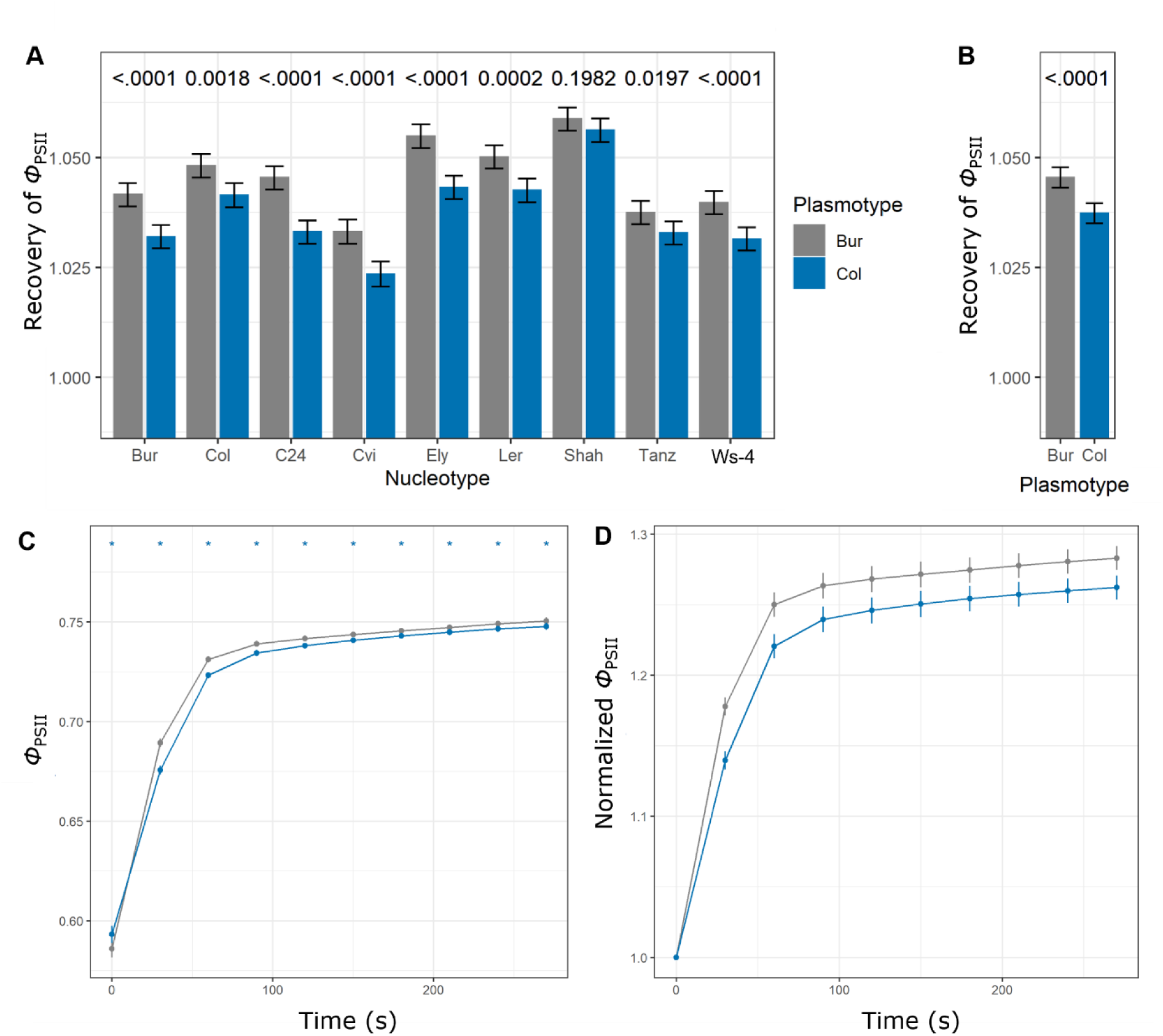
Bur-0 plasmotype has faster recovery of Φ_PSII_ after exposure to high-light. A) Recovery of Φ_PSII_ during the low-light period, between the Bur-0 and Col-0 plasmotype in nine different nuclear backgrounds. Cybrids from the spring 2022 experiment were cultivated in in semi-protected conditions, as shown in Figure 1A. B) Data as shown in panel A, but used to calculate the additive difference between the two plasmotypes. C) Independent experiment with cybrids grown in climate-controlled conditions, with the Bur-0, Col-0, C24 and Ler nucleotypes, to phenotype in the recovery of Φ_PSII_ during darkness, after high-light conditions (1000 µmol m^-2^ s^-1^). D) Same as panel C, but data are normalized to t=0 seconds per individual plant. In all cases the statistical difference, using a t-test (α = 0.05), is calculated between the plasmotypes, for panel A this means per nucleotype.

Using high-throughput chlorophyll fluorescence imaging (Figure 1B), amongst others to measure NPQ_(t)_ (as described by Tietz et al., 2017), we screened for plasmotypes with exceptional dynamic properties of NPQ. The cybrids were exposed to a six minute fluctuating light regime, alternating between low-light (200 µmol m^-2^ s^-1^) and high-light (1200 µmol m^-2^ s^-1^) (Figure 1C). Compared to the average of 60 plasmotypes, the Bur-0 plasmotype showed 23.0% lower NPQ_(t)_ 120 seconds after the transition to low-light from high- light (Supplementary Figure 3). The reduction in NPQ_(t)_ coincides with an increase of up to 2.1% in the light use efficiency of photosystem II electron transport (*Φ*_PSII_), which is also most pronounced during the recovery in low-light after a high-light period. This difference in NPQ_(t)_ and *Φ*_PSII_ is seen as an additive effect, as the reduction can be seen in all four nucleotypes. To assess whether this plasmotypic effect is additive in more than the four nucleotype backgrounds we generated a panel containing cybrids with the Bur-0 and Col-0 (as a control) plasmotypes and nine other globally representative nucleotypes. These cybrids where also grown in semi-protected conditions in two different seasons: autumn 2020 and spring 2021. While some phenotypes showed an epistatic relationship (i.e. interaction between the nucleotype and plasmotype), it was clear that the recovery of *Φ*_PSII_ was fastest in cybrids containing the Bur-0 plasmotype in all nucleotypes (Figure 2A and 2B). To reveal how the difference in dynamic responses of photosynthesis between the two plasmotypes behaved when allowed to recover for a longer period, four different nucleotypes combined with the Bur-0 and Col-0 plasmotypes were grown in controlled environments and exposed to fluctuating light. After 1000 µmol m^-2^ s^-1^, the irradiance was turned off, and the cybrids were phenotyped every 30 seconds. Here, *Φ*_PSII_ recovered faster and continued to be significantly higher due to the Bur-0 plasmotype (Figure 2C), an effect that is even more pronounced when we normalized to t=0 seconds (Figure 2D). Thus, within Bur-0 we identified allelic variation in the plasmotype for the relaxation of NPQ, which translates to a faster recovery of *Φ*_PSII_.

### Revealing allelic variation in the causal gene

It was then necessary to identify the allelic variant in the causal gene to gain more insights in the mechanisms that were responsible for this change in photosynthetic output. Due to the uniparental inheritance of the organelles, and thus the absence of recombination, mapping approaches commonly used to pin-point causal genes in the nuclear genome cannot be used. Within the cybrid panel we used, none of the other 59 plasmotypes possessed the faster recovery of *Φ*_PSII_ we observed with the Bur-0 plasmotype, and thus we can eliminate all those genetic variants that are shared between Bur-0 and any of the other plasmotypes. Using variant calling data, generated with a variant-calling pipeline customized for organellar genomes (Theeuwen et al., 2022a), we found that the Bur-0 has 33 and 8 SNPs and INDELs (i.e. maximum size 50bp) in the chloroplast genome and the mitochondrial genomes, respectively, that are absent in other plasmotypes (Supplementary Table 1). As this approach could not rule out larger structural variation causing the phenotypic difference, we made a *de novo* assembly of Oxford Nanopore Technology-derived large-read sequencing of the organellar genomes of Bur-0. This uncovered no structural variation in the chloroplast genome that went unnoticed in the whole genome resequencing dataset (Supplementary Figure 4B). However, the mitochondrial genome did show major rearrangements and three large copy number variants between Bur-0 and Col-0 (Supplementary Figure 4A). While such rearrangements are in line with the dynamic nature of the mitochondrial genome in plants (Mackenzie, 2007), these can result in the deletion of genes. In this case, none of the breakpoints are inside genes (Supplementary Table 2), and based on Illumina short read mapping we could see the same copy number variants also occur in plasmotypes that do not show the faster recovery of *Φ*_PSII_. Consequently, the phenotype can only be explained by SNPs or small INDELs, and for this reason we then predicted the effect of every variant. With this we found that only four SNPs are predicted to result in a non-synonymous change at the amino acid level (Supplementary Table 1). These four SNPs are spread over three genes, two in *MATURASE K* (*MATK*, referred to as SNPs *MATK^2084^* and *MATK^3108^*), one in *NAD(P)H-QUINONE OXIDOREDUCTASE SUBUNIT 6* (*NDHG*, referred to as SNP *NDHG^118888^*) and one in the chloroplast open reading frame 1 (*YCF1*, referred to as SNP *YCF1^124390^*).

To exclude more of the four non-synonymous SNPs, we used the variant calling data of 1531 publicly available accessions of *A. thaliana*, also generated with a variant-calling pipeline customized for organellar genomes (Theeuwen et al., 2022a). Amongst these we found three additional accessions, Cal-0, Cal-2 and NL1467, that possessed exactly the same four non-synonymous SNPs as Bur-0. Altogether the four accessions containing these SNPs were collected in Ireland, Great Britain and the Netherlands (Figure 3A), and all four accessions are genetically very closely related (Supplementary Figures 5, 6 and 7). Moreover, there are 117 accessions that cluster together with Bur-0, Cal-0, Cal-2 and NL1467 based on chloroplast genetic variation, and these possess two of the four remaining SNPs, namely the *MATK^2084^* and *YCF1^124390^*. In the absence of recombination in the chloroplasts, it is likely that mutations accumulate, implying there is likely to be an intermediate genotype where either the *MATK^3108^* or the *NDHG^118888^* SNP is present. As 20% of the tested accessions in the Netherlands have the *MATK^2084^* and *YCF1^124390^* SNPs, we screened an additional 1381 accessions collected from the Netherlands for the presence of the Bur-0 allele in either one of the *MATK* and *NDHG* genes (Figure 3B). This led to one accession, ID471, that was found to have the Bur-0 allele for *MATK^3108^*, but not for *NDHG^118888^*. Having accessions available that have fewer than the four SNPs that characterise the uniqueness of the Bur-0 plasmotype would allow us to pinpoint the causal mutation and thereby the causal gene.

**Figure 3.**
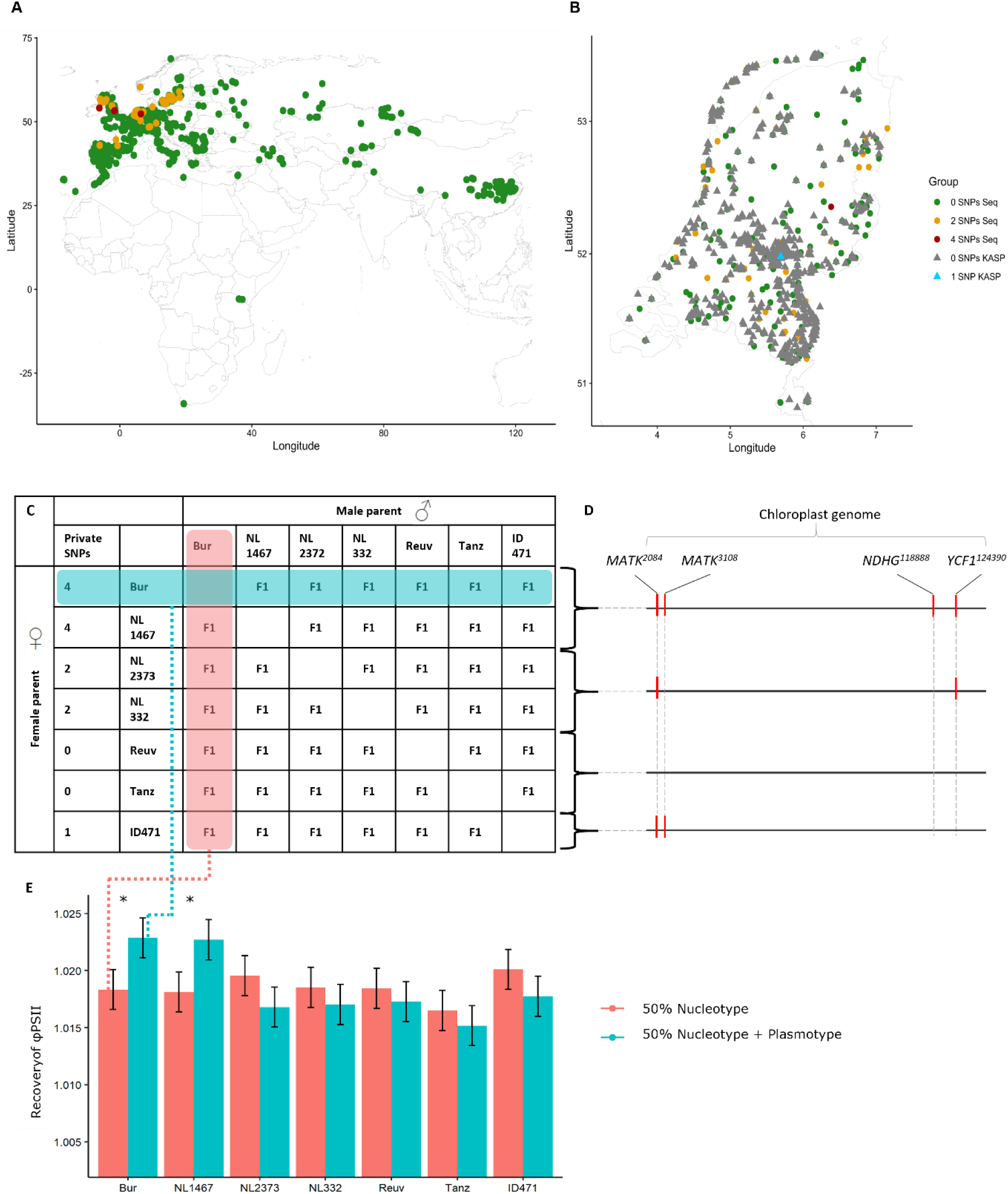
Genetic exclusion strategy to pinpoint NDHG as causal gene of the phenotypic difference. A) Distribution of alleles for MATK and NDHG in European, Asian and African continents. B) Distribution of alleles for MATK and NDHG within the Netherlands. C) Overview of the diallel panel used for this approach, F_1_ hybrids with the same nuclear combination were grouped according to the same plasmotype and all plasmotypes. D) Graphical representation of SNPs in the chloroplastic genome of the different accessions. E) Recovery of Φ_PSII_ in the different groups of F_1_ hybrids. A difference can either be a plasmotype or maternal effect.

To identify the causal SNP, we could make cybrids and assess the recovery of *Φ*_PSII_, and thereby link it to the causal SNPs, but this would require four generations of growing and crossing *A. thaliana*. Instead, we constructed a full diallel, entailing crossing accessions with different subset of the SNPs to generate reciprocal F_1_ hybrids (Figure 3C), a method only requiring one generation. The full diallel was made between two accessions with SNPs *MATK^2084^*, *MATK^3108^*, *NDHG^118888^* and *YCF1^124390^* (Bur-0 and NL1467), 2 accessions with SNPs *MATK^2084^* and *YCF1^124390^* (NL332 and NL2373), 1 accession with SNPs *MATK^2084^* and *MATK^3108^* (ID471) and 2 accessions with none of the private variants (Reuv-0 and Tanz-1)(Figure 3C and 3D). All F_1_ hybrids were phenotyped using the short fluctuating light protocol (Figure 1C), and recovery of *Φ*_PSII_ was measured. Next we compared the average recovery of *Φ*_PSII_ of the group of F_1_ hybrids having an accession as female parent (as depicted in blue in Figure 3C and 3E), with the reciprocal group of F_1_ hybrids having the same accession as the male parent (as depicted in red in Figure 3C and 3E), would reveal the plasmotypes showing the faster recovery of *Φ*_PSII_. This showed that Bur-0 and NL1467 were the only accessions showing a plasmotypic effect on recovery of *Φ*_PSII_, compared to the average of the plasmotypes (Figure 3E). As the other plasmotypes do not show the phenotypic effect, the SNPs in *MATK^2084^*, *MATK^3108^* and *YCF1^124390^* can be excluded, allowing us to pinpoint *NDHG* as the prime candidate gene causal to the phenotyping difference.

Since *NDHG* is a chloroplastic gene most transformation or gene editing methods cannot be used to test the allelic variants of *NDHG*. To find independent evidence that the cause of the faster recovery of *Φ*_PSII_ lay with variation in the *NDHG* alleles, we used specific cybrids that differed in the *NDHG* alleles to allow us to conduct more in-depth phenotyping. *NDHG* is part of the NAD(P)H dehydrogenase-like (NDH) complex, which redirects electrons from ferredoxin to the plastoquinone pool, while pumping protons from the stroma into the lumen; electrons from the plastoquinol pool can then be transferred via the cytochrome *b_6_f*, plastocyanin and the reaction centre of PSI to ferredoxin, thus forming a cyclic electron transport (CET) path around PSI (Shikanai, 2016; Strand et al., 2019). Within NDH, *NDHG* serves as one of the proton pumps, and an effect caused by *NDHG* should be measurable as a difference in the NDH activity measured via the post-illumination fluorescence rise (Shikanai et al., 1998). While this method previously has been used via non-imaging chlorophyll fluorometers, here we used the method as part of a high-throughput chlorophyll fluorescence imaging protocol. The use of knock-out (KO) mutants of two nuclear NDH genes, *ndho* and *ndhm,* that are in subcomplex A of the NDH complex, allowed us to see if the method is able to differentiate between differences in activity of the NDH complex. Comparing Col-0 with *ndho* and *ndhm* indeed shows the presence of a post-illumination fluorescence rise in Col-0 and its complete absence in the mutants, consistent with a substantial suppression of NDH activity (Supplementary Figure 8). From this we conclude that our method allows for the measurement of NDH activity. Cybrids containing the Bur- 0 plasmotype, in all four nucleotype backgrounds, show a significantly reduced NDH activity compared to the Col-0 plasmotype (Figure 4A and 4B). In addition, the nucleotypes also differ in their NDH activity, and strikingly the relative loss of NDH activity produced Bur-0 plasmotype compared to the Col-0 plasmotype is constant across the different nucleotypes (Figure 4B). This also means the *NDHG* allele in Bur-0 reduces the NDH activity, but does not completely disrupt its function, as is the case in the *ndho* and *ndhm* mutants. Due to the difference in NDH activity between the Col-0 and Bur-0 plasmotypes, together with the allelic variant in *NDHG* being the only option left to explain the observed phenotypic difference, we conclude that the *NDHG* allele causes the observed faster recovery of *Φ*_PSII_ associated with the Bur-0 plasmotype.

**Figure 4.**
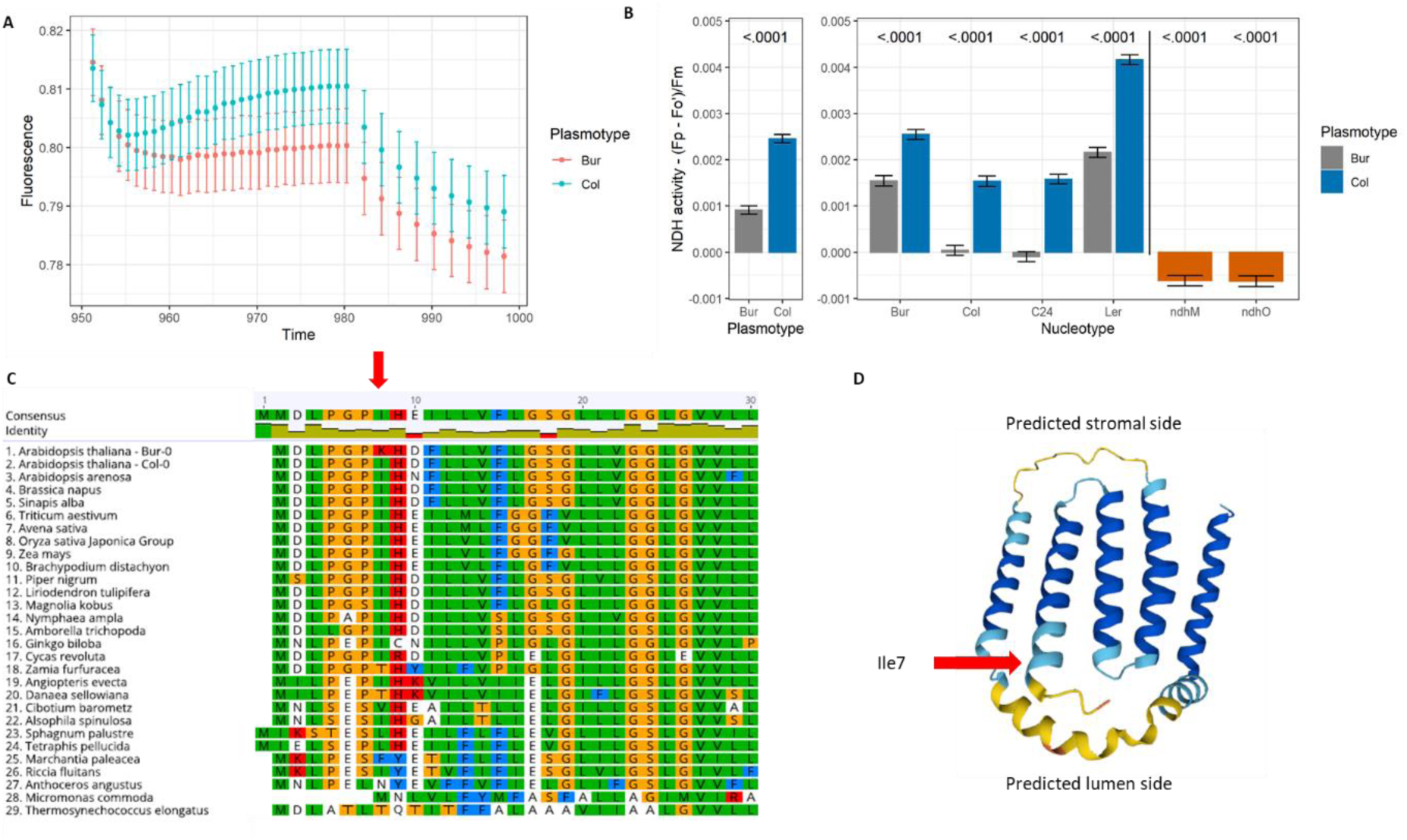
Differences in NDH activity and NDHG alleles between Bur-0 and Col-0 plasmotypes. A) Post illumination fluorescence rise during darkness, after 5 minutes of 200 µmol m^-2^ s^-1^. The data is normalized to the last measurement before darkness. Between 950 seconds and 980 seconds an F ’ measurement took place every second, while after 980 seconds the F’ measurement took place every two seconds, indicating a small effect of the measurement on the phenotype. B) Post illumination fluorescence rise as seen in panel A converted into NDH activity for the cybrids and ndho and ndhm mutants. C) Amino acid sequence alignment outlining the differences between the NDHG allele between Bur-0, Col-0 and more distantly related species. Ile7 or similar amino acids are highly conserved throughout the selected species apart from the exception Bur-0. The colours represent amino acid properties as defined by the Clustal colour scheme. D) Alphafold2 prediction of NDHG with the five transmembrane alpha helices. Ile7 is predicted to be located on the lumen side of the membrane.

In concluding that *NDHG* is the causal gene for the faster recovery of *Φ*_PSII_ in Bur-0 we sought to understand how the *NDHG* mutation in Bur-0 contributes to NDH activity. The *NDHG* allele in Bur-0, Cal-0, Cal-2 and NL1467 contains a Lys7, while all other *A. thaliana* accessions screened contain a Ile7. As these four accessions turned out to be very closely related both in nucleotype and plasmotype, it is likely that the Lys7 of *NDHG* only arose once in *A. thaliana*. Additional phylogenetic analyses reveal that Ile7 is highly conserved all the way back to the cycads and ferns, and even more ancient species contain amino acids in this position with similar properties to those of Ile (Figure 4C). The amino acid sequence at the N-terminal of *NDHG* is highly conserved, highlighting the biochemical importance and phenotypic effect of the substitution of the hydrophobic Ile7 to an positively charged Lys7 of *NDHG* in Bur-0. Ile7 is predicted to be in the first alpha helix on the lumen side of the NDH complex, possibly involved in proton release into the lumen (Figure 4D). Linking this with the reduced activity of the Bur-0 allele of *NDHG*, we conclude that the Lys7 in *NDHG* causes a reduction in the overall NDH activity.

### Natural variation to reveal insight into role NDH complex

To understand how the reduced NDH activity, as caused by the Bur-0 allele of *NDHG*, can be linked to faster recovery of *Φ*_PSII_ after a high to low-light transition, we further unravelled the consequences of the mutation. For this we reanalysed a dataset on cybrids, focussing on cybrids with the Bur-0 and Col-0 plasmotype in seven different nucleotypes grown in a high-throughput phenotyping system in which the light conditions can be dynamically controlled and all plants can be phenotyped simultaneously (Cruz et al., 2016; Flood & Theeuwen et al., 2020). The cybrids were grown for three weeks at 200 µmol m^-2^ s^-1^, and subsequently phenotyped at steady-state low-light (200 µmol m^-2^ s^-1^), sinusoidal high-light (up to 500 µmol m^-2^ s^-1^) and fluctuating light (up to 1000 µmol m^-2^ s^-1^) (Figure 5A). In this experiment we observed *Φ*_PSII_ and other PSII parameters to differ dynamically between the plasmotypes, depending on the light irradiance (Figure 5B). During steady-state low-light conditions *Φ*_PSII_ of Bur-0 was 1.1% higher than Col- 0, an effect that was independently confirmed in a separate high-throughput phenotyping system at steady-state low-light (200 µmol m^-2^ s^-1^) where especially after lights turned on *Φ*_PSII_ was 0.91% higher with the Bur-0 plasmotype (Figure 5C and 5D). During these steady-state low-light conditions we also observed a change in the ratio of the NPQ subcomponents in the presence of the Bur-0 plasmotype, where *q*_E_ is reduced by 5.2% and *q*_I_ is increased by 6.4% (Figure 5B). This effect on *Φ*_PSII_, *q*_E_ and *q*_I_ is largely absent during high-light either under steady-state or high-light periods in fluctuating light conditions, yet in the low-light periods in fluctuating light conditions the effects on *Φ*_PSII_, *q*_E_ and *q*_I_ are more pronounced compared to steady-state low-light. The largest difference is observed after 20 minutes at 191 µmol m^-2^ s^-1^ following 10 minutes at 500 µmol m^-2^ s^-1^, where *q*_E_ is then 16.9% lower and *Φ*_PSII_ is 1.8% higher in the presence of the Bur-0 plasmotype. Focussing on the responses to fluctuating light, and splitting the effects into high-light effects and low-light effects, we observed that during the low-light conditions the Bur-0 allelic variant of *NDHG* significantly influences *Φ*_PSII_, *q*_E_ and *q*_I_, and this effect is absent when light levels exceed ∼350 µmol m^-2^ s^-1^ (Figure 5D and 5E). Overall, we conclude that the allelic variant of *NDHG* in Bur- 0 causes a significant change in *Φ*_PSII_ and the components of NPQ during low-light conditions, and the effect is more pronounced when the cybrids are exposed to fluctuating light.

**Figure 5.**
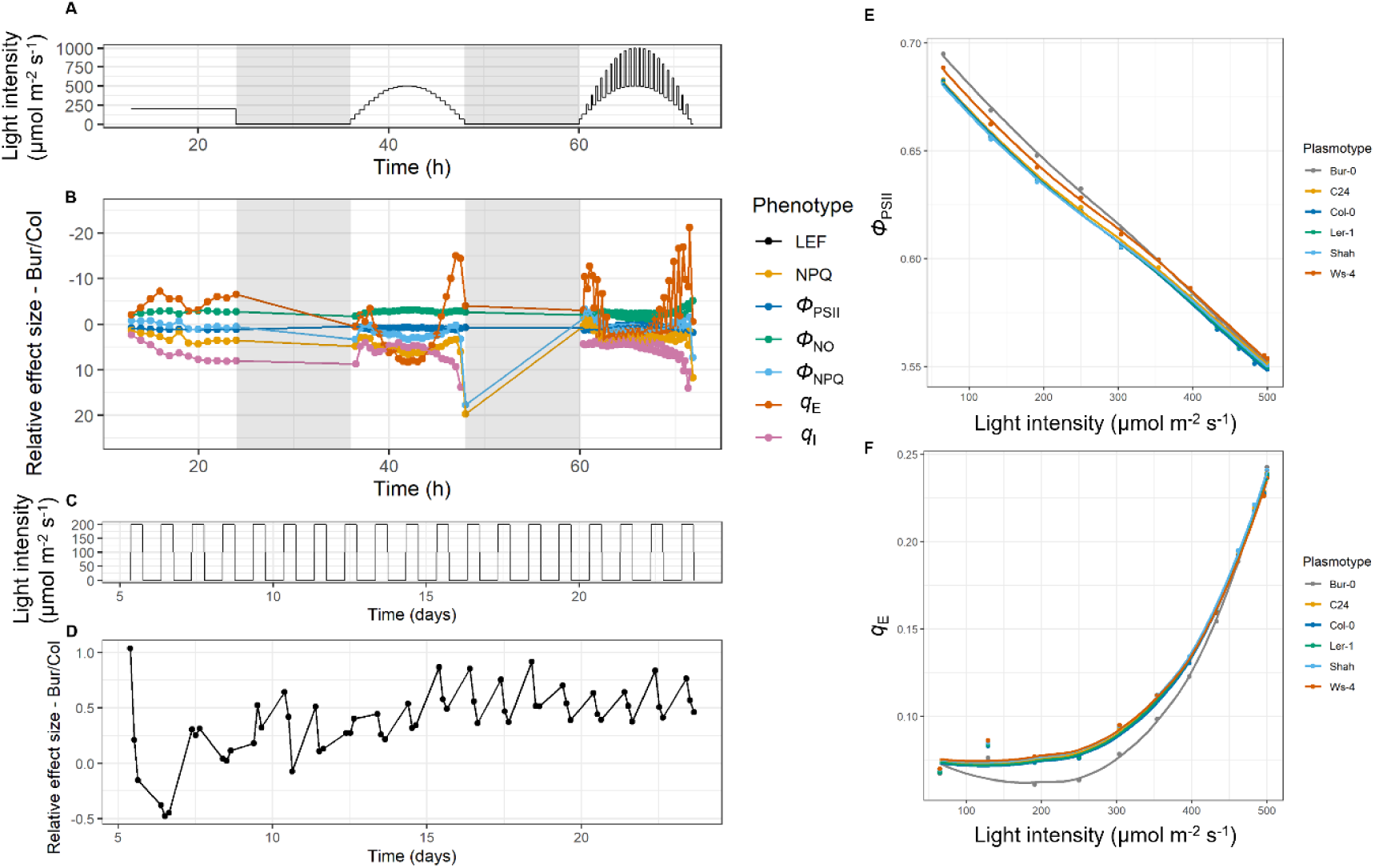
Differences in photosynthesis parameters under different light conditions. A) The light regime to which the cybrids were exposed after being grown for 21 days at 200 µmol m^-2^ s^-1^. The experiment was performed in the DEPI system (Cruz et al., 2016) B) The relative effect size between the Bur-0 and Col-0 plasmotypes for different photosynthetic parameters. C) The light regime to which the cybrids were exposed throughout the full experiment. D) The relative effect size between Bur-0 and Col-0 plasmotype for Φ_PSII_ as performed in the Phenovator system (Flood et al., 2016). E and F) Zoom in of the second half of the fluctuating light regime, and only the lower light conditions of the fluctuations. Points represent measurements after 20 minutes at the high to low-light transition.

The NDH complex is a key component of one of the CET pathways, in which, amongst others, it has a proton pumping capacity (Strand et al., 2017). This proton pumping capacity can influence the thylakoid lumen pH, which in turn is a major regulatory component in photosynthesis via the regulation of NPQ (Gilmore and Yamamoto, 1992). To reveal how the reduced NDH activity due to the Bur-0 plasmotype results in increased steady-state *Φ*_PSII_ in low-light conditions as well as recovery of *Φ*_PSII_ after fluctuations in light, we further examined the connection with NPQ. For this we phenotyped cybrids differing from the Bur-0 and Col-0 plasmotype in four nucleotype backgrounds, that adapted to fluctuating light conditions (20-minute fluctuations between 100 and 400 µmol m^-2^ s^-1^), every 30 seconds for 5 minutes at 1000 µmol m^-2^ s^-1^ and subsequently 5 minutes at 50 µmol m^-2^ s^-1^. During the high-light period no significant differences in *Φ*_PSII_ were observed, though in the low-light period *Φ*_PSII_ was significantly reduced for the duration of the 5 minutes, with the greatest difference appearing 30 seconds after the transition from high- light when *Φ*_PSII_ was 2.2% higher with the Bur-0 plasmotype (this is in line with relaxation in darkness as shown in Figure 1F)(Figure 6A). Interestingly, this increased *Φ*_PSII_ does not coincide with a faster relaxation of NPQ in all nuclear backgrounds (Figure 6D and Supplementary Figure 9). While the increased *Φ*_PSII_ is present in all nuclear backgrounds, ranging from 1.6 to 3.5% higher 30 seconds after high-light, the range of NPQ relaxation in the different nucleotypes is much wider (Supplementary Figure 9). In the Col-0 nuclear background NPQ is 1.9% lower, coinciding with an increase of 3.5% in *Φ*_PSII_, while in the C24 nuclear background NPQ is increased by 3.6%, but still *Φ*_PSII_ is increased with 2.2% (Supplementary Figure 9). The increase of *Φ*_PSII_ is compensated via a roughly equal division of *Φ*_NO_ and *Φ*_NPQ_, and thus no consistent difference on NPQ is observed (Figures 6B and C). As the Bur-0 *NDHG* allele causes a consistent increase in *Φ*_PSII_, but not a consistent reduction in NPQ, suggests that the increased *Φ*_PSII_ is not directly linked to faster NPQ relaxation.

**Figure 6.**
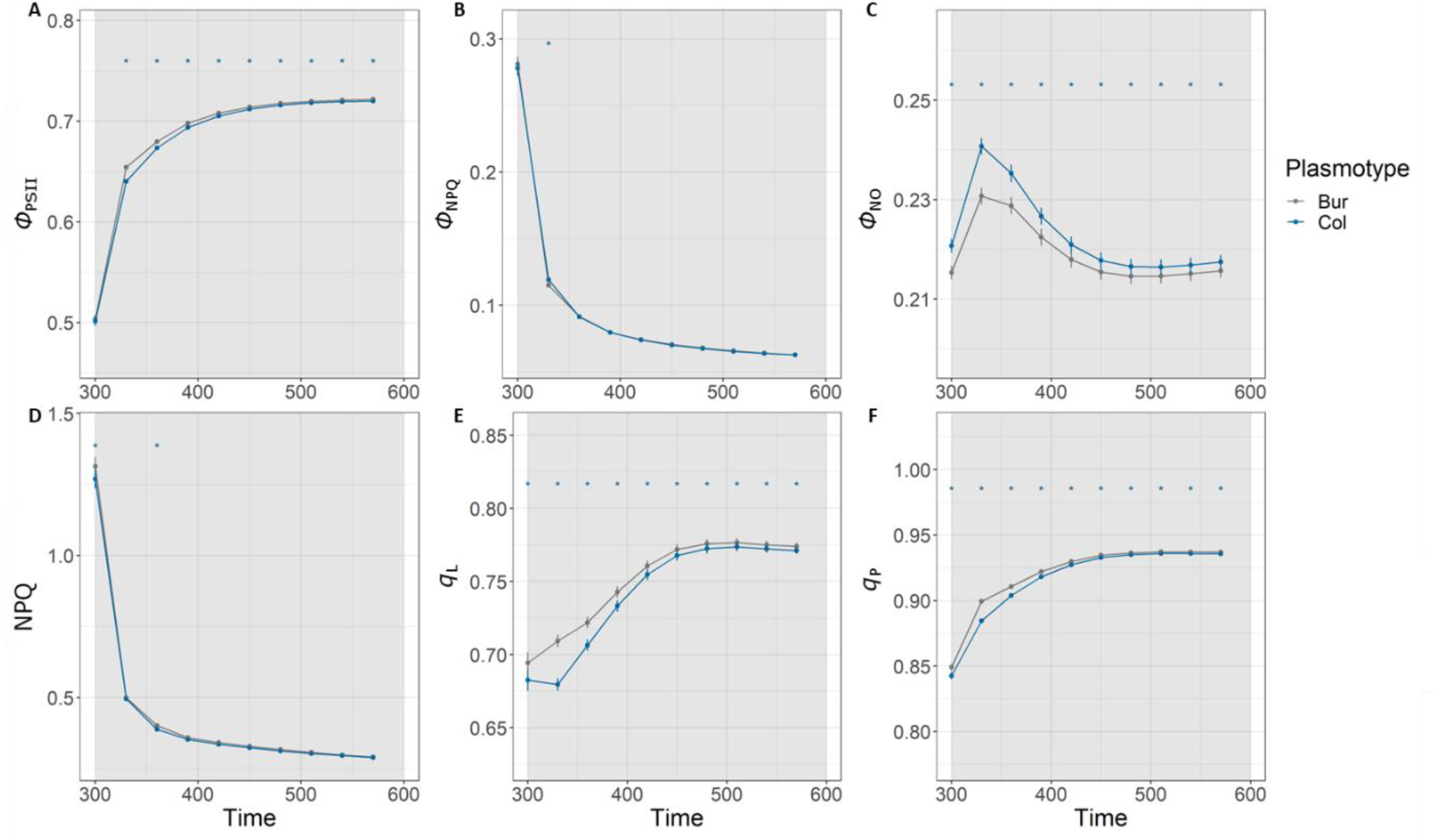
Phenotypic response of cybrids differing for the Bur-0 and Col-0 plasmotype during 50 µmol m^-2^ s^-1^ immediately following 5 minutes of 1000 µmol m^-2^ s^-1^. A – D show the additive effect of the two plasmotypes. See Supplementary Figures 9 and 10 for the response per individual cybrids, and the response of the ndho and ndhm mutants. Onset of 50 µmol m^-2^ s^-1^ is at t=300 seconds.

A reduction in NDH-mediated CET does not only result in the deposition of fewer protons in the lumen per unit linear electron transport (LET), but will also result in less reduction of the plastoquinone pool via the NDH CET route. Fewer electrons arriving into the plastoquinone pool via NDH could allow more LET, thus increasing *Φ*_PSII_ via an increase in the photochemical efficiency factor (*q*_P_) and the extent of open PSII reaction centres (*q*_L_). To this end we measured *q*_P_ and *q*_L_ simultaneously with the previously described *Φ*_PSII_ and NPQ. Here we observed the Bur-0 plasmotype to have on average 4.8% more open PSII reaction centres 30 seconds after a high-light treatment, which means a more oxidised Q_A_ pool and thus very likely a more oxidized plastoquinone pool; this coincides with 2.0% more photochemical quenching (Figure 6E and 6F). As with the higher *Φ*_PSII_ observed in all nuclear backgrounds, the effects on *q*_L_ and *q*_P_ are likewise present in all nuclear backgrounds (Supplementary Figure 9). Thus, while NDH mediated CET in some environmental conditions and nuclear backgrounds impacts NPQ, it more directly impacts the availability of the plastoquinone pool to accept electrons from PSII (and thus LET), allowing more efficient *Φ*_PSII_ and photosynthesis under conditions where ATP supply is not limiting.

### Is less NDH-mediated CET always beneficial?

While we found that lower NDH-mediated CET can result in a higher *Φ*_PSII_ under low-light conditions and faster recovery of *Φ*_PSII_ in fluctuating light conditions, the coinciding reduction in CET can have negative consequences as well. To assess the consequences of reduced NDH-mediated CET we phenotyped cybrids with both the Bur-0 and Col-0 plasmotypes in four nuclear backgrounds, as well as *ndho* and *ndhm* mutants in six different light regimes. These light regimes all had the same daylength, and three of the regimes had an average light intensity of 415 µmol m^-2^ s^-1^ (i.e. “Constant 415”, “Fluctuating 2 min” and “Fluctuating 10 min”), while the other three had an average irradiance of 340 µmol m^-2^ s^-1^ (i.e. “Constant 340”, “Fluctuating DEPI” and “Fluctuating maize”) (Figure 7A). The regimes ranged from steady-state light conditions (“Constant 415” and “Constant 340”), different regularly fluctuating light regimes (“Fluctuating 2 min” and “Fluctuating 10 min”), a sinusoidal fluctuating light regime (“Fluctuating DEPI”) and a highly fluctuating light regime as measured 1.5m above ground inside a mature maize crop on a windy day (“Fluctuating maize”)(Figure 7A). The plants were phenotyped using the 6-minute fluctuating light regime as used before (Figure 1C) and shoot dry weight was determined. When comparing the *ndho* and *ndhm* mutants against Col-0 60 seconds after high-light, *Φ*_PSII_ was significantly different between both NDH mutants and Col-0, in all six conditions. However, in four conditions *Φ*_PSII_ increased by on average 2.4% in the NDH mutants, but in NDH mutants exposed to the “Fluctuating 2 min” and “Fluctuating maize” regimes, *Φ*_PSII_ was reduced with 2.4% and 12.7%, respectively, implying that the absence of NDH mediated CET can have a positive impact on *Φ*_PSII_, but only if fluctuations in light are not too frequent (Figure 7B). Despite this, in all regimes except the “Fluctuating 10 min” (here *ndho* is found to have 16.3% more shoot dry weight compared to Col-0), at least one of the two NDH mutants shows a reduction in shoot dry weight.

**Figure 7.**
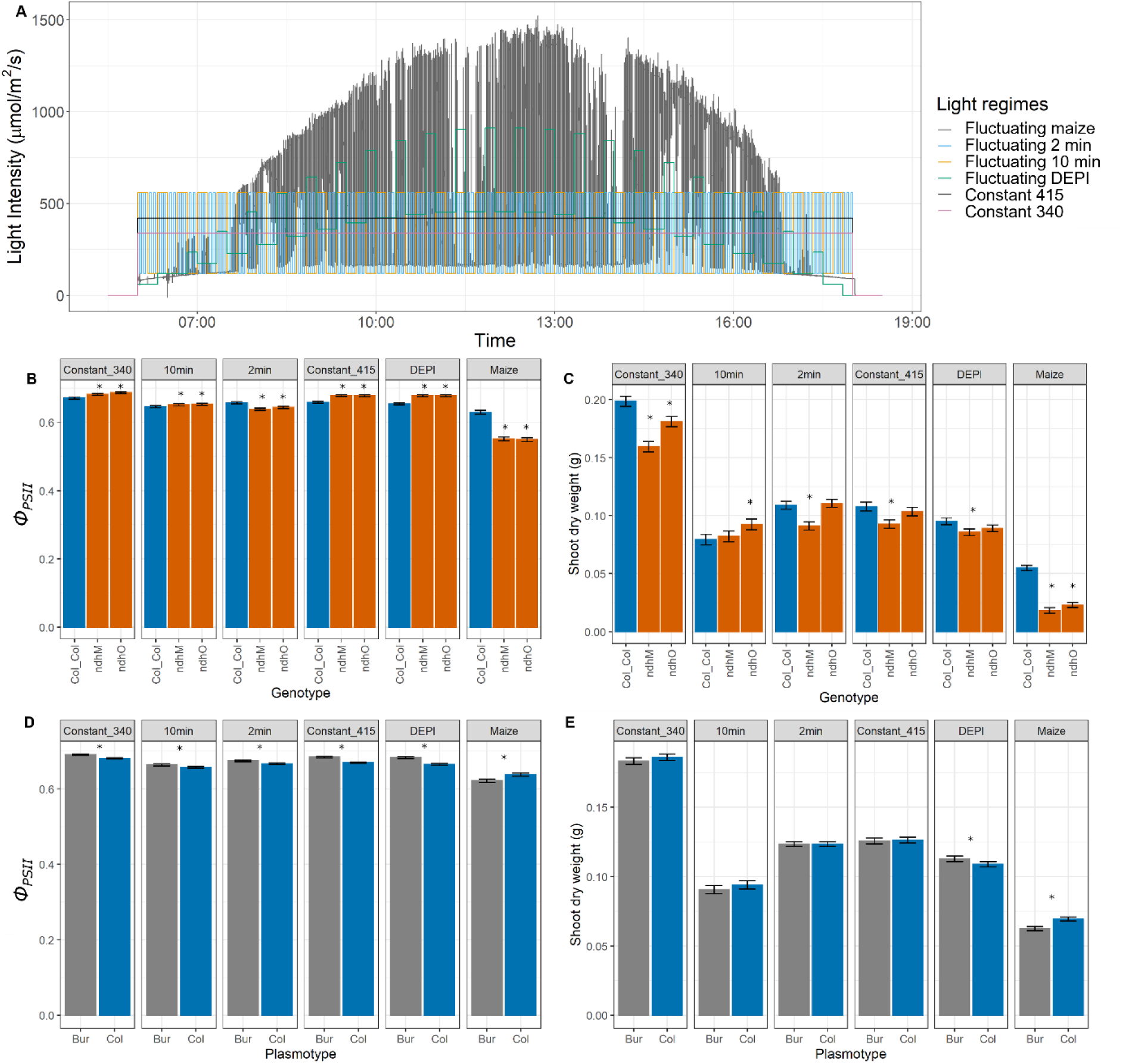
Assessing the trade-off between Φ_PSII_ and NDH-mediated CET in different light regimes. A) The six different light regimes as the plants were exposed. B) The effect of Φ_PSII_ between Col- 0 and ndho and ndhm mutants in the different light regimes. C) The effect on shoot dry weight between Col-0 and ndho and ndhm mutants in the different light regimes. D) The additive effect of Φ_PSII_ between Col-0 and Bur-0 plasmotypes in the different light regimes. E) The additive effect on shoot dry weight between Col-0 and Bur-0 plasmotypes in the different light regimes. Note that the experiments were performed in two separate experiments, with “Constant 340”, “Fluctuating DEPI” and “Fluctuating maize” together in one experiment, and “Constant 415”, “Fluctuating 2 min” and “Fluctuating 10 min” together in another experiment. The first three regimes had an average light irradiance of 340 µmol m^-2^ s^-1^, and chlorophyll fluorescence-based phenotyping was done on 25 DAS and shoot dry weight was harvest at 26 DAS. The other three regimes had an average light irradiance of 415 µmol m^-2^ s^-1^, and chlorophyll fluorescence-based phenotyping was done on 26 DAS and shoot dry weight was harvest at 30 DAS.

As a result of the “Fluctuating maize” regime, where *Φ*_PSII_ showed the greatest reductions, an average reduction in shoot dry weight of 62.7% when comparing the *ndho* and *ndhm* mutants against Col-0 was observed (Figure 7C). This shows that while a seemingly positive impact on *Φ*_PSII_, due to reduced NDH mediated CET, most light regimes tested still results in a reduced shoot dry weight. This illustrates a clear trade-off between *Φ*_PSII_ and NDH-mediated CET, that in five out of the six light regimes tested results in reduction in shoot dry weight.

The absence of a positive effect on shoot dry weight in the NDH mutants that show an increased *Φ*_PSII_ in less stressful conditions might either be due to the absence of CET outweighing the increased *Φ*_PSII_ or simply that there are additional bottlenecks limiting the translation of increased *Φ*_PSII_ into increased shoot dry weight. As the allelic variant of *NDHG* in Bur-0 has a less reduced NDH-mediated CET as compared to the NDH mutants in conditions where less CET is required, the benefit of increased *Φ*_PSII_ might actually benefit plant growth. Indeed, while the NDH mutants had a reduced *Φ*_PSII_ of 2.4% 60 seconds after a high to low-light light transition, when grown in the “Fluctuating 2 min” regime the Bur-0 plasmotype had a 1.1% increase (Figure 7B and 7D). Likewise, in the “Fluctuating maize” regime *Φ*_PSII_ was 2.6% lower in the Bur-0 plasmotype, in comparison to the 12.7% reduction in the NDH mutants. In “Fluctuating maize” this resulted in a 10.1% reduction of shoot dry weight as conveyed by the cybrids with the Bur-0 plasmotype, compared to the Col-0 plasmotype. As the reduction due to the NDH mutants was 62.7% in comparison to the Col-0 wildtype, this indicates that in these conditions the reduced NDH-mediated CET, as a result of the *NDHG* allele of Bur-0, also has less of an reducing effect on shoot dry weight. So, while CET is less impacted in the *NDHG* allele of Bur-0, in the four conditions (“Constant 415”, “Constant 340”, “Fluctuating 10 min” and “Fluctuating DEPI”) where NDH mutants had an average 2.4% increase in *Φ*_PSII_, the Bur-0 plasmotype still had a 1.8% increase in *Φ*_PSII_. Besides a significant 3.6% increase in shoot dry weight in cybrids with the Bur-0 plasmotype, as compared to the Col-0 plasmotype, when exposed to sinusoidal fluctuating light conditions (“Fluctuating DEPI”), we did not observe consistent positive effects on shoot dry weight as a result of the *NDHG* allele of Bur-0. This might be the result of NDH-mediated CET playing a bigger role than described, also in less stressful conditions (i.e. fluctuating light conditions), but that the benefit of increased *Φ*_PSII_ masks the effect of reduced or downregulated CET. Taken together we conclude that there are conditions where both the NDH mutants as well as the *NDHG* allele of Bur-0 convey a negative effect on plant growth, and in one case a small significant positive effect.

## Discussion

Using a species-representative cybrid panel (Theeuwen et al., 2022a), grown in semi-protected cultivation conditions, we screened for divergent NPQ relaxation characteristics due to natural genetic variation in the plasmotypes. Besides differences in NPQ, we observed a faster recovery of *Φ*_PSII_ after a high to low-light transition whenever the Bur-0 plasmotype was present (Figure 2). This effect seemed to be robust because it was observed in spring and autumn 2020 and spring 2021 even though all three growing seasons presented differing climatic trends. Previously, the Bur-0 plasmotype had been shown to produce a small, but significant, effect on the photosynthetic phenotypes (Flood & Theeuwen et al., 2020). The experimental setup used in Flood & Theeuwen et al., 2020 only allowed for a measurement of *Φ*_PSII_ 20 minutes after a high to low-light transition while our protocol examined the changes in *Φ*_PSII_ during the first 90 seconds after the high to low irradiance transitions. This revealed a more pronounced impact of the Bur-0 plasmotype on the dynamics of *Φ*_PSII_. Consistent with Flood & Theeuwen et al., 2020 that the effect size of this trait was largest in fluctuating light, we conclude that the effect of the Bur-0 plasmotype was primarily associated with the fast responses to dynamic light conditions. *q*_E_ (here measured as NPQ) is a measure of the activation or capacity of the photoprotective mechanism, and is not a measure of the yield of dissipation of PSII excited states of chlorophyll via the *q*_E_ quenching mechanism, a proxy for the *Φ*_NPQ_ parameter (Kramer et al., 2004). Therefore, the finding of faster recovery of *Φ*_PSII_ conveyed by the Bur-0 plasmotype to be associated with faster relaxation of *Φ*_NPQ_ implies there is not merely a difference in activation of NPQ but also in actual dissipation. Hence the variation causal to the faster recovery of *Φ*_PSII_ is promising for potential photosynthetic improvements, as has been experimentally shown (Kromdijk et al., 2016; De Souza et al., 2022).

One way to unravel the underlying physiological basis of this phenotypic difference would be to identify the underlying genetic basis. However, due to the uniparental inheritance of the plasmotype and consequential absence of recombination, standard approaches to mapping the genes underlying phenotypic variation are not possible, neither could we resort to transformation approaches. Although recent developments in organellar transformation technology are promising, so far these methods will neither work for the adenine to thymine (or vice versa) nucleotide change nor the substitution of the native *NdhG* allele (Ruf et al., 2019; Kwak et al., 2019; Jakubiec et al., 2021; Kang et al., 2021; Nakazato et al., 2021). Instead, we used a full diallel consisting of reciprocal hybrids that differ only for their plasmotype, which allows to determine whether a given plasmotype conveyed the faster recovery of *Φ*_PSII_ or not. By including accessions differing for the remaining candidate SNPs in the full diallel, a genetic exclusion approach can be used. However, a genetic exclusion approach relies on the genetic variation available within a population. In using the 1,531 accessions for which publicly available sequencing information was available, we found accessions that would allow the narrow down two candidate genes to explain the phenotypic difference. We were only able to make the genetic exclusion approach work when we found an extra accession that differed in one of the two remaining candidate genes in 1,381 accessions from the Netherlands that were additionally screened. The use of these accessions in the full diallel revealed allelic variation for *NdhG* to be associated with the faster recovery of *Φ*_PSII_ (Figure 3). Further confirmation of *NdhG* being the causal gene relied on a high-throughput chlorophyll fluorescence imaging protocol to measure NDH activity via the post-illumination fluorescence rise. Previously, this method had exclusively been used via non-imaging chlorophyll fluorometers to measure chlororespiration, a process involving NDH (Feild et al., 1998), and later, more explicitly, NDH activity (Shikanai et al., 1998). Nuclear NDH mutants validated the approach in our phenotyping system and consequently revealed a consistent difference in NDH activity between the Bur-0 and Col-0 plasmotypes in combination with different nucleotypes (Figure 4). This led us to conclude the Ile7 to Lys7 substitution in *NdhG* is causal to the increase in recovery of *Φ*_PSII_. Furthermore, we also found substantial variation in NDH activity due to the nucleotype, a source of variation that should be further examination.

While the NDH complex is arguably dispensable, and even shown to be absent in some lineages of orchids and the gymnosperms, the fact that 29 genes persist in most plants families underlines the evolutionary constraint on the NDH complex and alleles thereof (Lin et al., 2017; Strand et al., 2019). Whereas in *A. thaliana* so far no conditions were found where NDH mutants played a significant role in plant performance (Munekage et al., 2004; Okegawa et al., 2008; Suorsa et al., 2012), while in rice the NDH complex is shown to play an important role in fluctuating light conditions (Yamori et al., 2016). We show a reduced shoot biomass of 62.7% in the *ndho* and *ndhm* mutants when subjected to highly fluctuating light conditions (Figure 7) in contrast to the Col-0 control, demonstrating the importance of the NDH complex. The mutation in the Bur-0 allele of *NdhG* is likely to have an impact on NDH is suggested by the fact that the N-terminus of *NdhG* is highly conserved, with the Ile7 being conserved since the common ancestor with the cycads and ferns. This was found to be the case in highly fluctuating light conditions where the Bur-0 allele of *NdhG* reduces shoot biomass with 10.1%, in contrast to the Col-0 allele, suggesting the conservation is due to otherwise significant impacts on plant performance. This is contradictory, while the finding that the Ile7 to Lys7 substitution results in phenotypic differences, an RNA editing mechanism was shown to induce a Ser17 to Phe17 substitution, but was not found to impact the functioning of the NDH complex (Okuda et al., 2010). Overall, these observations leads to two (apparently contradictory) conclusions; on one hand, some species seem to have lost genes encoding for the NDH complex, and there is substantial nuclear variation for NDH activity suggesting that NDH is ancillary. On the other hand, the highly conserved *NdhG* gene, including the Ile7 amino acid, and the findings that the Ile7 to Lys7 conversion and the NDH mutants result in substantially impacted plant performance suggest that it is essential in these species (e.g., via its role in cyclic electron transport).

As the NDH complex is a key part of one of the CET pathways, variance in NDH activity could allow differential control of photosynthesis via the ratio between the formation of reducing power and the transthylakoid proton motive force. The latter is the driving force for ATP synthesis and consists of two components, the transthylakoid voltage or potential and the transthylakoid proton concentration (or pH) difference. It can also act to decrease lumen pH, an element of proton motive force. CET as a whole can therefore, for example, vary the ratio of NADPH and ATP production in the light dependent reactions of photosynthesis and decrease lumen pH. In the case of NDH mediated CET, reduced ferrodoxin, produced on the acceptor side of PSI, is used by NDH to reduce plastoquinone. This process releases four protons per electron into the thylakoid lumen (Strand et al., 2017). Oxidation of plastoquinol by the cytochrome *b_6_f* complex, a process ultimately dependent on PSI photochemistry, releases another two protons per electron into the lumen. The thylakoid lumen pH, a component of the transthylakoid pH difference is important in controlling the rate constant of the electron transfer step between plastoquinol and the cytochrome *b_6_f* complex, and thus rate of thylakoid electron transport, and also the formation of zeaxanthin and protonation of *PsbS*, both of which lead to increased *q*_E_ (Kono et al., 2014). Lumen pH is therefore a major regulatory component in photosynthesis. As demonstrated the reduced NDH mediated CET is the result of allelic variation in the NdhG gene, which is found to be a component of the four proton pumps of the NDH complex (Strand et al., 2017), we extrapolate that the reduced NDH activity results in less protons being deposited in the thylakoid lumen. Using this rational, reduced NDH mediated CET would result in higher lumen pH, and thus result in less *q*_E_ (and therefore NPQ), which in turn would allow for increased LET and the observed increased recovery of *Φ*_PSII_. Nonetheless, while in all nuclear backgrounds a consistent faster recovery of *Φ*_PSII_ was observed, this did not consistently correlate with faster NPQ relaxation, implying the reduced proton pumping capacity cannot explain the observed phenomena. The absence of a diminished *q*_E_ following the high to low irradiance transition in all nuclear backgrounds means there is additional nuclear encoded variation for amongst others lumen pH regulation and antenna complexes, but also that there must be a more direct link between NDH mediated CET and PSII that explains in more rapid recovery of *Φ*_PSII_ associated with the Bur-0 plasmotype.

Increased *Φ*_PSII_ can be the result of increases in the photochemical efficiency factor (*q*_P_) and the extent of open PSII reaction centre (*q*_L_) (Kramer et al., 2004; Baker, 2008). *q*_P_ is non-linearly related to the redox state of Q_A_, the primary stable electron acceptor of PSII, and *q*_L_ is similar to *q*_P_ but adjusted so that it is a linear measure of Q_A_ redox state and thus the extent of open PSII reaction centres. As the post illumination fluorescence rise, the metric used to quantify NDH activity *in vivo*, is the result of reduction of the plastoquinone pool (Shikanai et al., 1998), NDH activity implicitly is an indication of electrons being deposited into the plastoquinone pool. Therefore, besides reduced NDH mediated CET due to the Bur-0 plasmotype, which results in the deposition of fewer protons per electron in the lumen, simultaneously the NDH complex deposits fewer electrons into the plastoquinone pool. The resulting more oxidised Q_A_ pool will result in a higher *q*_P_ and therefore PSII efficiency, enhancing LET. The increase in *q*_P_ (and *q*_L_) was most pronounced in the first 90 seconds after the high to low-light transition and consistently correlated with increases in *Φ*_PSII_ in every nuclear background (Figure 6). Despite the absence of a consistent impact on *q*_E_, variation in NDH activity is found to be directly related to the regulation of PSII via *q*_P_ and the plastoquinone pool. Furthermore, here we observed this difference in NDH activity, due to the Bur-0 plasmotype, to be most pronounced in the low light periods during fluctuating light and the effect disappears when irradiance exceeds 350 µmol m^-2^ s^-1^ (Figure 5). This observation would be in line with the hypothesis that the NDH complex plays a relatively small role in photosynthesis and a reduction in the NDH activity will only have an effect on photosynthetic performance in relatively low-light conditions, and, as shown here, especially during fluctuating light conditions (Burrows et al., 1998; Shikanai et al., 1998; Munekage et al., 2004).

Despite NDH-mediated CET primarily impacting photosynthesis in low light conditions, the observed physiological consequences of allelic variation in *NdhG* alongside the effects of the knock-out mutants of *ndho* and *ndhm*, suggests the NDH complex plays a substantial role in overall plant performance (as measured in terms of shoot biomass; Figure 7). However, as faster recovery of *Φ*_PSII_ in dynamic light conditions is shown to be able to contribute to increases in yield (Kromdijk et al., 2016; De Souza et al., 2022), we set out to test whether the potential benefit of faster recovery of *Φ*_PSII_ can outweigh to negative consequences as a result of reduced NDH-mediated CET. As this balance might potentially be different depending on the environmental conditions, we measured shoot biomass for cybrids with four different nucleotypes and either the Bur-0 or Col-0 plasmotypes in light regimes that primarily differed in the frequency of light fluctuations. In the regimes with constant light and fluctuations every 2 and 10 minutes, we observed no significant differences in shoot biomass, while in all these conditions the Bur-0 plasmotype shows to have the ability of faster recovery in *Φ*_PSII_ (Figure 7). Conversely, when the cybrids were grown in highly fluctuating light conditions, as measured inside a maize canopy on a windy day, the cybrids with the Bur-0 plasmotype are found to have reduced recovery of *Φ*_PSII_. In these conditions shoot biomass due to the Bur-0 plasmotype is reduced by 10.1% compared to the Col-0 plasmotype. Remarkably, in fluctuating light conditions following a sinusoidal pattern during the photoperiod, shoot biomass due to the Bur-0 plasmotype is increased by 3.6%. Why the Bur-0 allele of *NdhG* is beneficial in these conditions remains to be seen, but recent evidence suggests that sinusoidal light regimes differentially impact shoot biomass in comparison to square wave light regimes (Schiphorst et al., unpublished). So, while the Bur-0 allele of *NdhG* conveyed increased recovery of *Φ*_PSII_ with a potentially beneficial impact on yield via its effect on *q*_P_, the coincident reduction NDH mediated CET appears to negatively affect yield, revealing a trade-off between two effects and for which the outcome depends on the environmental conditions.

The trade-off between the speed of *Φ*_PSII_ recovery and NDH-mediated CET seems favourable, or is at least neutral, to retain NDH mediated CET at wildtype Col-0 levels in most environmental conditions. This would be in line with the observation that the Bur-0 *NdhG* allele is rare amongst *A. thaliana* accessions. Besides the *NdhG* allele in Bur-0 (sampled in Ireland), it was only found to be present in accessions Cal-0, Cal-2 (both sampled in Great Britain) and NL1467 (sampled in the Netherlands). All four accessions turned out to be genetically almost identical in both nucleotype and plasmotype. While the absence of the same plasmotype, or same SNP but different plasmotype, elsewhere in the world can be coincidental, the conservation of Ile7 across the species tree and within *A. thaliana* implies conditions that the phenotype arising from the Ile7 for Lys7 substitution is less fit. We found the *NdhG* allele of Bur-0 impacted overall growth the least in low and slowly fluctuating light conditions, which coincides with the fact that Ireland, England and the Netherlands are amongst the countries with the least sun hours year round within the species range of *A. thaliana* (Suri et al., 2020). In addition, we observed that the overall photosynthetic response of the plasmotype depends on the nuclear background, suggesting the nuclear background also plays an important role. Strikingly, this is in agreement with the observation that the Bur-0 accession was amoung the top ten with highest *Φ*_PSII_ out of 674 global accessions grown and phenotyped in steady-state low-light conditions (Flood, 2015). Moreover, within the Netherlands, NL1467 (regarded as nearly identical to Bur-0) was the best performing accession in terms of *Φ*_PSII_ out of a collection of 192 accessions (Wijfjes et al., unpublished). While in both cases the Bur-0 and NL1467 had the *NdhG* allele that conveyed up to a 1.1% higher *Φ*_PSII_ in the same low-light conditions (Figure 5), this cannot entirely explain the observed high *Φ*_PSII_, implying that the nucleus possess genetic variation for higher *Φ*_PSII_ as well. Due to this higher *Φ*_PSII_ in the nucleus, the reduction in plant performance as caused by the *NdhG* allele in the plasmotype, might be partially compensated for, to an extent in which the Bur-0 *NdhG* allele would be able to persist. Together with the native environmental conditions were the Bur-0 *NdhG* allele is found being neutral or even beneficial, this could explain how the Ile7 to Lys7 substitution could occur and be conserved.

Here, the first systematic exploration of dynamic photosynthetic properties due to natural variation in the organellar genomes, revealed that carrying a mutant allele of *NdhG* led to faster recovery of *Φ*_PSII_ after a high to low irradiance step. While a faster recovery of *Φ*_PSII_ has previously been shown to cause positive effects on yield, here we show that the faster recovery of *Φ*_PSII_ is directly caused by a reduction in NDH mediated CET. While previous studies in *A. thaliana* suggest that the NDH complex plays an insignificant role in photosynthetic performance, we conclude the contrary. Based on our results, we believe there is a trade-off between the potentially faster recovery of *Φ*_PSII_ and the negative consequences arising from the reduction in NDH mediated CET. Further studies into the physiological properties of the trade-off will reveal whether there is potential to recruit the beneficial properties on recovery of *Φ*_PSII_ without the negative consequences of NDH-mediated CET, possibly by enhancing other CET pathways. However, it is interesting why other CET pathways cannot compensate for the relatively small effect in NDH-mediated CET as caused by the Bur-0 allele of *NdhG*, and there are similar effects observed due to the nuclear genetic variation.

This might even suggest another mechanistic role for *NdhG* or the NDH complex as a whole. In conclusion, here we show how the plasmotype represents a source of untapped photosynthetic variation for which the genetic variation is worth being used, and for which the phenotypic consequences may hold a wealth of novel physiological insights.

## Material and Methods

### Plant material

The 232 cybrids from the cybrid panel are constructed and genotyped by Theeuwen et al., 2022a, were used in this work. Ten additional cybrids, where the Col-0 and Bur-0 plasmotype was combined with C24, Ler-1, Sha, Ws-4 and Ely nucleotypes, were used as constructed and described in Flood & Theeuwen et al., 2020. Dutch accessions NL1467, NL2371, NL332 and Reuv were obtained from Wijfjes et al. (unpublished). The additional 1,381 Dutch accessions screened in this paper, including ID471, were collected within the Netherlands, as described by Wijfjes et al. (unpublished) (also see “Acknowledgements”). The full diallel was made by crossing Bur-0, NL1467, NL2371, NL332, Reuv and Tanz reciprocally, the crosses for ID471 x Reuv failed and thus were excluded from the experiments. NDH mutants *ndho* (N568922) and *ndhm* (N587707) were obtained from the Nothingham Arabidopsis Stock Centre. The mutants were checked for homozygosity of the T-DNA insertion, where *ndho* was found to be homozygous for the T-DNA insertion, and for *ndhm* a heterozygous plant was selfed and amongst the offspring a plant that was homozygous for the T-DNA insertion was selected (Supplementary Table 4).

### Sequence analysis

#### Genetic exclusion approach

The genetic exclusion approach was based on variant calling data obtained from raw paired-end sequencing data as publicly available for 1,531 accessions (Theeuwen et al., 2022a). See Theeuwen et al., 2022a for the custom pipeline with filtering steps for ensuring high quality variants for the chloroplast and mitochondrial genomes. The predicted impact of private SNPs and INDELs of the Bur-0 plasmotype, in comparison to the other plasmotypes within the cybrid panel, was done using SnpEff (Cingolani et al., 2012). With the most recent version of the *A. thaliana* reference genome from TAIR10.1 (Sloan et al., 2018), also which contains an updated mitochondrial genome, a SnpEff database was constructed using a GFF genome annotation file obtained from the NCBI taxonomy ID 3702 page. Variants with “HIGH” and “MODERATE” impacts were considered to have a potential impact on gene function. The genetic relationship between accession, based on nuclear, mitochondrial and chloroplast variant calling data, was determined using a hierarchical clustering analysis the vcfR package (version 1.13.0) and the dist.gene and as.phylo functions from the ape package (version 5.6-2).

#### De novo assembly

As input material for the *de novo* assembly of Bur-0 wildtype, 500 mg of young leaf material was used. High-quality high molecular weight (HMW) DNA was extracted following the LeafGo protocol (Driguez et al., 2021). Using 100 ng/µL HMW DNA (Qubit dsDNA Quantitation BR Kit) short reads were eliminated with the PacBio SRE Circulomics Kit. 1 µg HMW DNA (Qubit dsDNA Quantitation BR Kit) was used to do end-prepping and nick repairing using the NEBNext Companion Module Kit (#E7180S), followed by ligation using the Oxford Nanopore Technologies (ONT) Ligation Sequencing Kit (SQK-LSK109). Roughly 50X coverage, after basecalling, was generated using an ONT Flow Cell (R9.4.1) on the MinION Mk1C. The basecalling was done using the “fast basecalling” option the on the MinION Mk1C. For the nuclear genome all reads we assembled using the Flye assembler, using default settings (Kolmogorov et al., 2019). However, this did not result in continuous assemblies of the organellar genomes. Instead, we separately extracted all reads that mapped to the chloroplast or mitochondrial TAIR10.1 reference genomes using BLAST. Next, for the chloroplast reads we filtered out 250 reads larger than 50 kbp, and for the mitochondrial reads we filtered out 1,500 reads larger than 20 kbp. The chloroplast and mitochondria were assembled using the Flye assembler, resulting in continuous assemblies of the organellar genomes. The contigs where polished in four subsequent rounds using Pilon in default settings (Walker et al., 2014) with the short read Illumina data generated for the Bur-0 wildtype in (Flood & Theeuwen et al., 2020). Synteny analysis of the chloroplast and mitochondrial genomes was done using SyRi (Goel et al., 2019), and analysis of the breakpoints of the mitochondrial genome was done using Geneious Prime. The annotated gene file of TAIR10.1 was used to determine whether genes were located on the breakpoints.

#### Dutch population screening

Plants were grown in controlled environmental greenhouse conditions, and 1 cm^2^ leaf material was collected in 96 deepwell plates, and DNA was extracted using a SPRI Bead Technology-based DNA extraction protocol. Using KASP markers (Supplementary Table 4) for SNPs at *MATK^3108^* and *NDHG^118888^*, the alleles for all 1,381 Dutch accessions the were determined. Mapping of accessions was done using rworldmap (version 1.3-6), rworldxtra (version 1.01), mapproj (version 1.2.8) and ggplot2 (version 3.3.6).

#### NDHG amino acid analysis

BLASTP was used to collect the *NDHG* gene of the species as named in Figure 3C, and Geneious Prime was used to align the genes. The predicted structure of *A. thaliana NDHG* was retrieved from AlphaFold2 via UniProt.

### Phenotyping experiments and data analysis

#### Semi-protected conditions experiments

Plant growth took place in an outdoor, gauze covered tunnel at Unifarm, Wageningen University and Research, the Netherlands (51.9882583, 5.66119897). The base of the tunnel measured 8 × 5 m and was enclosed by synthetic gauze material that was largely penetrable by rainwater, sunlight and wind so to provide conditions similar to those encountered in the field. Rain gauges placed inside and outside the tunnel confirmed that all rainwater was able to penetrate the gauze. Light irradiance was measured both inside the tunnel and at a metrological station within 500 m from the tunnel. A comparison of readings from both locations, during the growing period of spring 2020, indicated that the gauze decreased the light irradiance that penetrated through to the growing area on average by 94.3 µmol m^-2^ s^-1^. The floor of the tunnel was covered in black landscape material and the tunnel contained a zipper door to prevent the entry of any undesired interferents.

Black plastic pots measuring 7×7×18 cm (Bestebreurtje B.V., Huissen, the Netherlands) were used for individual plants as they allowed sufficient depth for unlimited root development. Grey plastic trays measuring 40×60×20 cm were used to hold 40 of the aforementioned pots. The trays were organized in five rows of 11 trays and one row of 13 trays. A small seedling tray was placed in the bottom of each large grey tray to raise the pots above the edge of the grey tray. The plant pots were filled with a mixture of 40 % sand and 60 % peat provided by Lensli Substrates (Katwijk, the Netherlands). The substrate includes YARA PG MIX™ which contains 15-10-20+3 of N, P_2_O_5_, K_2_O and MgO. The added fertilizer is in powder form and results in complete substrate values of 1.0 and 5.7 for electrical conductivity and pH, respectively. No additional nutrition was applied during the experiment.

In spring of 2020 plastic wrapping was placed over the trays during the night for the first 14 days of growth to protect from cold temperatures. The same was done in spring 2021 for the first five days of growth and in autumn 2020 this was not done at all. In spring and autumn 2020 grey rubber covers were placed on each pot after seedlings had established, leaving 0.5 cm space for water to reach the soil. In spring 2021 the soil was covered with blue Friedola Mega Stop mats as described in (Junker et al., 2015) and provided by TEUN B.V. (Rucphen, the Netherlands). The pots were evenly watered as needed according to weather conditions and rainfall. Anti-slug/snail pellets were placed in small piles on the ground around the perimeter of the tunnel. Remote sensors from 30MHz (Amsterdam, the Netherlands) monitored and recorded light irradiance and temperature every minute at plant level.

Phenotyping was primarily done using the high-throughput phenotyping system PlantScreen^TM^ SC System provided by Photon Systems Instruments spol. s r.o (Drásov, Czech Republic). Using the chlorophyll fluorescence camera and a RGB camera, we obtained a range of photosynthetic and morphological parameters. A custom made 6-minute fluctuating light protocol (Figure 1C), with 5 F’ and Fm’ measurements, allowed us to calculate *Φ*_PSII,_ *Φ*_NOt,_ *Φ*_NPQt_, NPQt and *q*_E_ at different moments during the 6- minute protocol. A custom R script converted these measurements into 37 parameters capturing the dynamic response to the fluctuating light protocol. A custom mask using the Schedular software was generated to phenotype 20 plants, with a 7 x 7 cm grid size. In spring and autumn 2020, the grey rubber plates were complemented with additional rubber strips, to mask soil-grown algae being registered. In spring 2021 the blue mats ensured the algal growth was not recorded. Automatic masking by the Data Analyser software ensured background noise was removed, and the plant mask was generated. The RGB camera generated an additional 20 parameters quantifying the morphological characteristics and colour properties. In spring 2020 the shoots were also harvested to measure the shoot dry weight when the first flower of a plant opened.

During spring 2020 221 cybrids were ready and could be sown. An R script was used to create an unbalanced, incomplete block design to randomize the cybrids among the pots resulting in the nucleotypes being randomized among the trays and the plasmotypes being randomized within the trays. The number of replicates ranged between 10-12 for the cybrid genotypes and 60-80 for the four WTs. Due to the number of pots, sowing was spread over three days, with cybrids with the Bur-0 nucleotype being sown on March 18^th^ 2020, cybrids with Col-0 and Cvi nucleotypes on March 19^th^ and cybrids with Tanz nucleotypes on March 20^th^. Approximately 4 pre-germinated seedlings were sown per pot, and all but the healthiest seedling was removed 20 days after sowing. The trays containing nucleotypes Cvi-0, Col-0, Bur-0 and Tanz-1 were phenotyped in the PlantScreen^TM^ system at 35, 38, 38 and 38 days of growth, respectively.

During spring 2021 all 240 cybrids were ready and could be sown. A complete randomized block design was used, in a way that all 240 cybrids were sown once every six trays. Such six trays were arranged in a 2 x 3 orientation, and together formed one big block, with the total experiment having n=12. As phenotyping in the PlantScreen^TM^ system allowed only 20 plants, we ensured that all cybrids within a block were measured back-to-back to correct over the day and time as best as possible. Again, the sowing was split over three days, cybrids with the Bur-0 nucleotype were sown on March 17^th^ 2021, cybrids with the Col-0 nucleotype on March 18^th^, and cybrids with the Cvi and Tanz nucleotype on March 19^th^. After 24 days of growth, all but the healthiest seedling were removed. Phenotyping in the PlantScreen^TM^ system happened on May 3^rd^ and 4^th^, meaning the plants ranged between 45 and 48 days old.

Data from the two consecutive spring experiments were analysed and combined. As shown previously (Theeuwen et al., 2022a), eight cybrids turned out to not have the desired genotype, and thus were excluded in all data analysis steps. Observations on plants that either did not germinate or established badly, in contrast to the other replicates, were removed if 1.5 standard deviation smaller than the average of a genotype per year. Here we were interested in additive plasmotype effects, and thus we ignored any interaction effects (those are further analysed in Theeuwen et al., 2022a). To calculate the Best Linear Unbiased Estimators, we used a linear mixed model via lme4 (version 1.1-30), where each block comprised the combination of blocks within each year. The equation used:

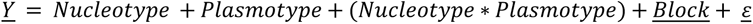

During autumn 2020 a subset of cybrids was grown, where cybrids with the Col-0 and Bur-0 plasmotype was combined with nine different nucleotypes, and the Col-0 and Bur-0 wildtypes grown as controls. The plants were grown in a complete randomized block design, consisting of 12 blocks, each with 40 plants (and thus compromise one tray), totalling n=24. All seedlings were sown on October 2^nd^ 2020 and 27 days later, all but one plant was removed. 75 days after sowing all plants were phenotyped in the PlantScreen^TM^ system, and 88 days after sowing the shoots were harvested to determine shoot dry weight. Observations on plants that either did not germinate or established poorly, in contrast to the other replicates, were removed if 1.5 standard deviation was smaller than the average of a genotype. Next all parameters were analysed using a linear mixed model approach, using the restricted maximum likelihood procedure with the lme4 package (version 1.1-30). Using analysis of variance with the Kenward-Roger approximation for degrees of freedom, significant differences were calculated between the Col-0 and Bur-0 plasmotypes, within one nucleotype and overall nucleotypes, with a threshold in the post hoc tests of α = 0.05. The linear mixed model equation also included the X and Y coordinates, as some edges showed faster drying out, compared to the middle:

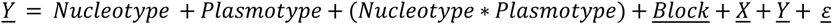

#### DEPI experiment

This experiment was based on reanalysed data from Flood & Theeuwen et al., 2020. Here, the two separate experiments, both including the full 7 x 7 cybrid panel, were analysed together, resulting n=8 replicates. We only analysed the first 3 days, as one of the two experiments only had data for the first 3 days. While we were interested in the additive effect differences between the Col-0 and Bur-0 plasmotype only, we ran a linear mixed model with the equation:

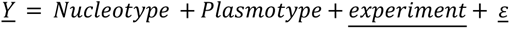

including all cybrids, to allow estimating the experiment effect best. Subsequently, we extracted the pairwise differences between the Col-0 and Bur-0 plasmotypes for all parameters. For the fluctuating light analysis (Figure 5E and 5F) we extracted the parameters as measured in the low light periods, on the second half of the third day.

#### Phenovator experiment

For this experiment we reanalysed data as published in Flood & Theeuwen et al., 2020. The analysis was identical as describe above in the “DEPI experiment”, but instead there was data for n=24 and 25 days in total.

#### NPQ relaxation and NDH activity experiment

Cybrids and the NDH mutants were grown in a climate chamber with 10-hour photoperiod and a light irradiance of 250 µmol m^-2^ s^-1^, for the first 19 days after sowing. Next the plants were exposed to fluctuating light irradiances, alternating between 100 and 400 µmol m^-2^ s^-1^ every 20 minutes, during the 10-hour photoperiod. During the photoperiod the temperature was 20 °C and during the night 18 °C, and throughout the experiment the relative humidity was kept at 70%. Plants were grown on a 4 x 4 x 4 cm rockwool substrate provided by Grodan B.V. (Roermond, the Netherlands), and irrigated weekly with a nutrient solution (Supplementary Table 3). Plants were sown in a complete randomized block design, with n=24. After 24 days of growth, plants were phenotyped in the PlantScreen^TM^ system. Here all plants were dark adapted for 30 minutes, to retrieve Fo and Fm. Next plants were exposed to five minutes of 20 µmol m^-2^ s^-1^, and during the subsequent 50 seconds the post illumination fluorescence rise was monitored in darkness, with the first 30 seconds a measurement every one second, and the remaining 20 seconds a measurement every 2 seconds. This was repeated with the only difference that the five minutes were set at a light irradiance of 200 µmol m^-2^ s^-1^. Following this, the six minutes fluctuating light protocol was, as shown in Figure 1C, with fluctuations between 200 and 1000 µmol m^-2^ s^-1^. Then for five minutes plants were exposed to 1000 µmol m^-2^ s^-1^. Lastly, for five minutes, the lights were turned off, and F’ and Fm’ were measured every 30 seconds. Next all parameters were analyzed using a linear mixed model approach, using the restricted maximum likelihood procedure with the lme4 package (version 1.1-30). Using analysis of variance with the Kenward-Roger approximation for degrees of freedom, significant differences were calculated with a threshold in the post hoc tests of α = 0.05. In this experiment we tested for significant differences between the Col-0 and the two NDH mutants, using the following equation:

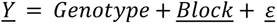

and for significant additive differences between the Col-0 and Bur-0 plasmotypes, using the following equation:

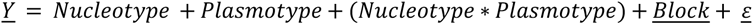

#### Full-diallel experiment

The full diallel was grown in a climate chamber with 10-hour photoperiod and a light irradiance of 250 µmol m^-2^ s^-1^, for the first 16 days after sowing. Next the plants were exposed to fluctuating light irradiances, alternating between 100 and 400 µmol m^-2^ s^-1^ every 20 minutes, during the 10-hour photoperiod. During the photoperiod the temperature was 20 °C and during the night 18 °C, and throughout the experiment the relative humidity was kept at 70%. Plants were grown on a 4 x 4 x 4 cm rockwool substrate provided by Grodan B.V. (Roermond, the Netherlands), and irrigated weekly with a nutrient solution (Supplementary Table 3).

Plants were sown in a complete randomized block design, with n=12. On day 23 after sowing, all plants were phenotyping the PlantScreen^TM^ system, using the 6-minute fluctuating light regime, yielding 37 chlorophyll fluorescence and 20 morphological parameters. Outliers were removed when either not germinated or badly established, using a R script to remove plants two standard deviations smaller than the average per genotype per treatment, on the basis of plant leaf area. The raw kinetic data was converted into the photosynthetic parameters using a R script. Next all parameters were analysed using a linear mixed model approach, using the restricted maximum likelihood procedure with the lme4 package (version 1.1-30). Best Linear Unbiased Estimates were calculated using a model in which each F_1_ hybrid was used twice, once when belonging to the group with the same maternal parental genotype, and once when belonging to the group with the same paternal parental genotype. The linear mixed model used for this is as follows:

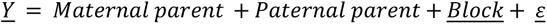

#### qP recovery experiment

Cybrids and the NDH mutants were grown in a climate chamber with 10-hour photoperiod and a light irradiance of 250 µmol m^-2^ s^-1^, for the first 14 days after sowing. Next the plants were exposed to fluctuating light irradiances, alternating between 100 and 400 µmol m^-2^ s^-1^ every 20 minutes, during the 10-hour photoperiod. During the photoperiod the temperature was 20 °C and during the night 18 °C, and throughout the experiment the relative humidity was kept at 70%. Plants were grown on a 4 x 4 x 4 cm rockwool substrate provided by Grodan B.V. (Roermond, the Netherlands), and irrigated weekly with a nutrient solution (Supplementary Table 3). Plants were sown in a complete randomized block design, with n=24. After 24 days of growth, plants were phenotyped in the PlantScreen^TM^ system. Here all plants were dark adapted for 30 minutes, to retrieve Fo and Fm. Next were exposed to 10 minutes of 50 µmol m^-2^ s^-1^ without measurements. The five minutes of 1000 µmol m^-2^ s^-1^ and five minutes of 50 µmol m^-2^ s^-1^ were started. During each of the five minutes, F’ and Fm’ were measured at the start (t=0s) and every 30 seconds afterwards. In between each 30 seconds measurements, 11 seconds of light irradiance (either 1000 or 50 µmol m^-2^ s^-1^) was given, after which six seconds of far-red was applied, followed with two seconds of darkness to calculate Fo’. The remaining 11 seconds was again set at the given light irradiance. Plants 1.5 standard deviations smaller than the average plant leaf area per genotype were removed. The raw kinetic values were converted to aforementioned photosynthetic parameters, but specifically for this experiment Fo’ was used to calculate *q*_P_ and *q*_L_ (Maxwell and Johnson, 2000). Next all parameters were analysed using a linear mixed model approach, using the restricted maximum likelihood procedure with the lme4 package (version 1.1-30). Using analysis of variance with the Kenward-Roger approximation for degrees of freedom, significant differences were calculated with a threshold in the post hoc tests of α = 0.05. In this experiment we tested for significant differences between the Col-0 and the two NDH mutants, using the following equation:

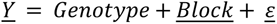

and for significant additive differences between the Col-0 and Bur-0 plasmotypes, using the following equation:

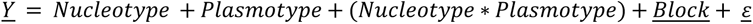

#### Biomass experiments in different light regimes

The cybrids and NDH mutants were grown in a climate chamber, with three light treatments at once, so split over two experiments. In both experiments the photoperiod was 12 hours, and during the photoperiod the temperature was 20 °C and during the night 18 °C, and throughout the experiment the relative humidity was kept at 70%. Plants were sown in a complete randomized block design, with two replicates per block, and 12 blocks per treatment (n=24). Plants were grown on a 4 x 4 x 4 cm rockwool substrate provided by Grodan B.V. (Roermond, the Netherlands), and irrigated weekly with a nutrient solution (Supplementary Table 3). All plants were phenotyped in the PlantScreen^TM^ system, using the six-minute fluctuating light regime, yielding 37 chlorophyll fluorescence and 20 morphological parameters. For the first experiment the phenotyping took place 25 days after sowing, and the shoot harvest 26 days after sowing. For the second experiment the phenotyping took place 26 days after sowing, and the shoot harvest 30 days after sowing.

In the first experiment the three light regimes had an average light irradiance of 340 µmol m^-2^ s^-1^, the “Constant 340” received this as a steady-state light condition and the “Fluctuating DEPI” received this in a sinusoidal fluctuating light regime inspired by the third day of the DEPI experiment as described above. The “Fluctuating maize” received this light as replicated to mimic measurements on September 20^th^ 2020 in Wageningen (The Netherlands, 51°59’20.0“N 5°39’43.2”E) 1.5 m above ground in a mature maize canopy. The light intensity dataset was measured using a quantum sensor (Licor) coupled to a transimpedance amplifier and recorded every 100 ms with a datalogger (ADC-24, Picolog). The light irradiance and daylength were transformed to get to a photoperiod of 12 hours and 340 µmol m^-2^ s^-1^ irradiance. In the second experiment the three light regimes received an average light irradiance of 415 µmol m^-2^ s^-1^, with the “Constant 415” receiving this as steady-state light, the “Fluctuating two min” and “Fluctuating 10 min” receiving this in periods of two and 10 min respectively while alternating between 120 to 560 µmol m^-2^ s^-1^. To generate a homogeneous light area (less than 5% variation over the growing area), we used 5 Fluence VYPR 2p (Austin, United States of America) per treatment. The light intensity was dynamically controlled using a custom-built digital dimmer. The digital dimmer consisted of a microcontroller (ESP32) interfaced to the light setup through an optocoupler which provided a variable 0- 10V dimming signal.

Outliers were removed when either not germinated or badly established, using a R script to remove plants 1.5 standard deviations smaller than the average per genotype per treatment, on the basis of plant leaf area. Next all parameters were analysed using a linear mixed model approach, using the restricted maximum likelihood procedure with the lme4 package (version 1.1-30). Using analysis of variance with the Kenward-Roger approximation for degrees of freedom, significant differences were calculated with a threshold in the post hoc tests of α = 0.05. In this experiment we tested for significant differences between the Col-0 and the two NDH mutants, using the following equation:

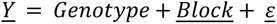

and for significant additive differences between the Col-0 and Bur-0 plasmotypes, using the following equation:

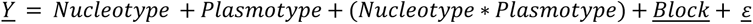

## Data availability

Both the raw Oxford Nanopore Technology sequencing data and the *de novo* assembly of Bur-0 will be available via the European Nucleotide Archive. Raw data will be available via supplementary datasets to this paper.

## Code availability

Measuring protocols as used on the Fluorcam7 software, R scripts for downstream analyses and the *de novo* sequence assembly pipeline of the Bur-0 genome will be available from GitLab (https://git.wur.nl/tom.theeuwen/bur-ndh-project).

## Acknowledgements

The authors would like to thank Maarten Koornneef for the insightful discussions throughout the whole project and feedback to the manuscript. Furthermore, we acknowledge Saskia van Dijk, Anne-Fleur Peters and Marco Albiero for their help in experimental work, Corrie Hanhart, Gema Flores Andaluz, René Boesten, Sanne Put, Louise Logie, Delfi Dorussen and Hedayat Bagheri for their help in sowing for different experiments, Raúl Wijfjes for his advice on *de novo* sequencing assemblies. We are also thankful of Joost Keurentjes who, together with EW, set up an initiative via the Dutch radio program “Vroege Vogels” (https://www.bnnvara.nl/vroegevogels) to have listeners collect Dutch *A. thaliana* seeds, and all the people who did so.

## Author contributions

TPJMT, JH and MGMA conceived and designed the study with substantial input by DMK and EW. TPJMT, AWL, DT, FF, FFMB and NF performed experiments and analysed data while the project was ongoing, TPJMT analysed the final data as presented in this paper. LC designed the custom-made light controller and TPJMT and LC build and optimized the system. TPJMT wrote the manuscript, with significant contributions by DMK, JH and MGMA. All authors read and approved the final version of the manuscript.

## Supplementary Tables

**Supplementary Table 1.**
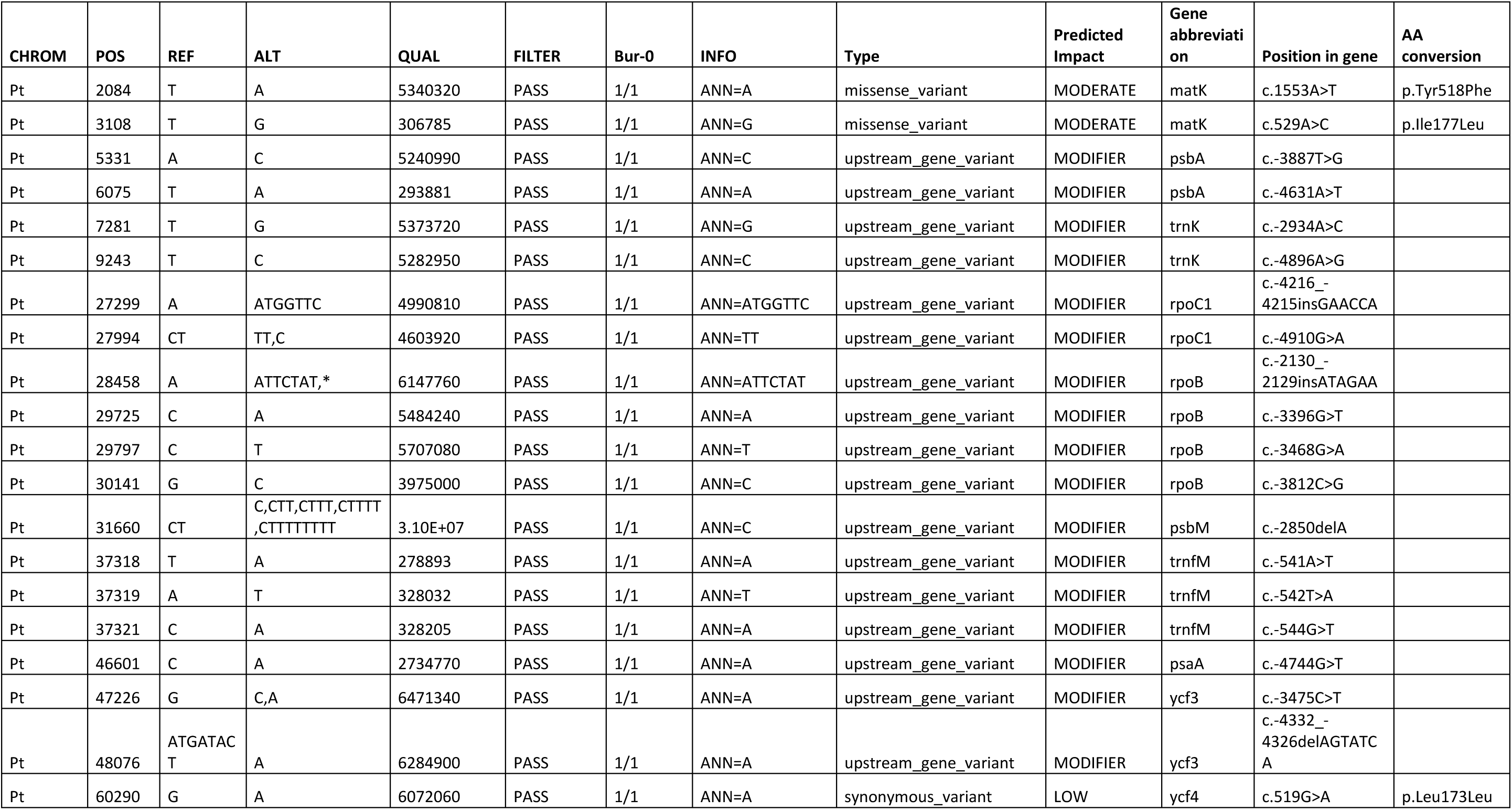
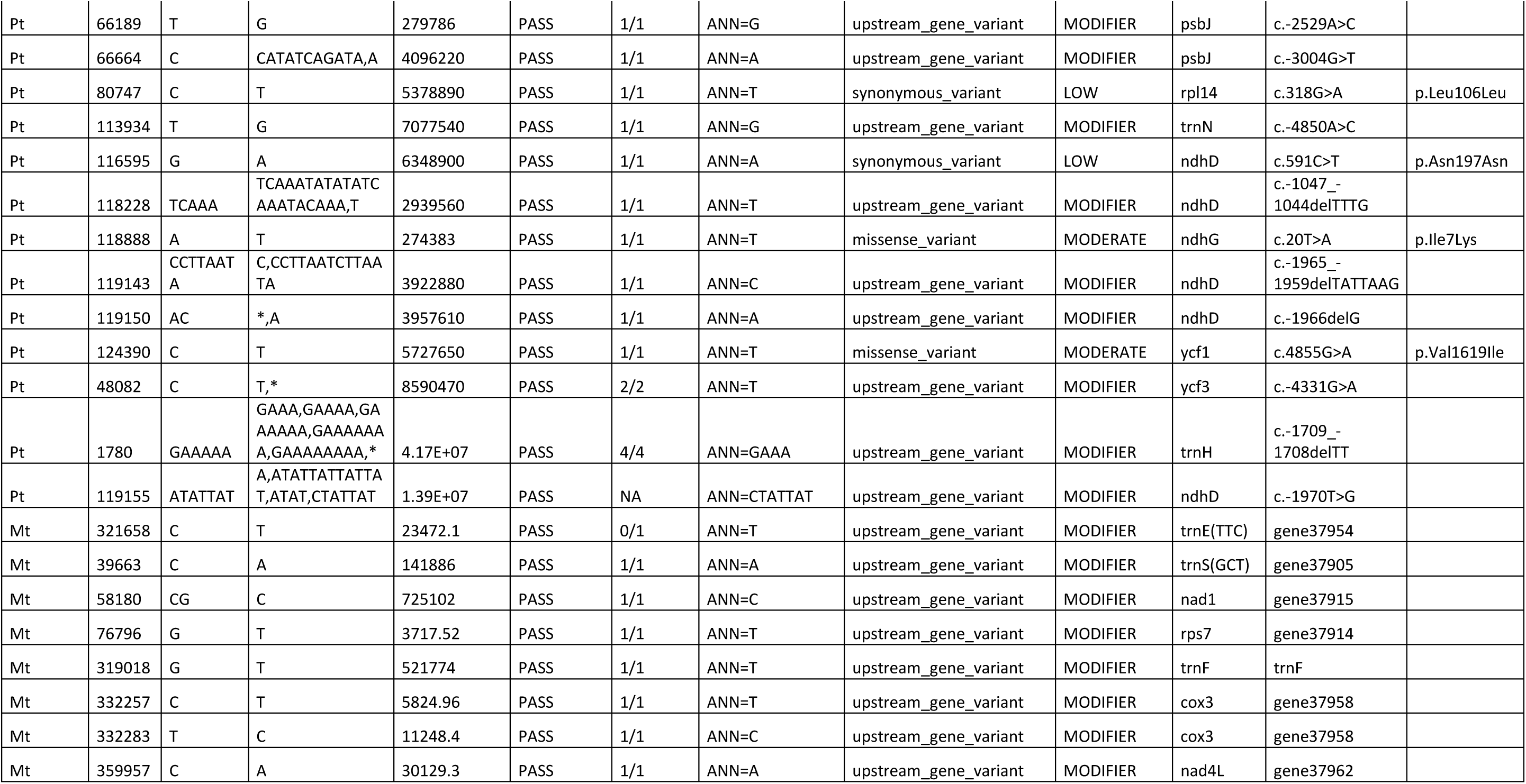
Overview of plasmotypic variants, as found unique in Bur-0 in contrast to the other accessions that are part of the cybrid panel. The position is based on the reference genome TAIR10.1. The variant prediction is based on SnpEff.

**Supplementary Table 2.** Breakpoints in alignment between Col-0 and Bur-0. The “Start” and “End” columns represent at which position the breakpoint starts and ends, compared to the Col-0 reference genome. The “Length” column represents the length of the breakpoint, and the “Result” column defines whether this region is a duplication, deletion or insertion. In the “Predicted impact” column the predicated annotation of the genome is used to determine what the impact of the breakpoint is.

| Start of breakpoint | End of breakpoint | Length (bp) | Result | Predicted impact |
| --- | --- | --- | --- | --- |
| 4192 | 4194 | 2 | Deletion | Intergenic |
| 6118 | 6570 | 452 | Duplication | cox2, but due to overlap the entire copy is present once |
| 30723 | 31348 | 625 | Deletion | Intergenic |
| 36362 | 36819 | 457 | Duplication | Intergenic |
| 41464 | 41997 | 533 | Duplication | Intergenic |
| 84089 | 84541 | 452 | Duplication | Intergenic |
| 143953 | 144410 | 457 | Duplication | Intergenic |
| 255123 | 270775 | 15652 | Deletion | atp6.2 is deleted |
| 277634 | 277670 | 36 | Deletion | Intergenic |
| 321435 | 321968 | 533 | Duplication | Intergenic |
| 332126 | 332181 | 55 | Deletion | Intergenic |

**Supplementary Table 3.** Nutrient solution as used for growing A. thaliana on rockwool substrate.

|  | Concentration | Unit |
| --- | --- | --- |
| NH<sub>4</sub> | 1.7 | mmol/L |
| K | 4.13 | mmol/L |
| Ca | 1.97 | mmol/L |
| Mg | 1.24 | mmol/L |
| NO<sub>3</sub> | 4.14 | mmol/L |
| SO<sub>4</sub> | 3.14 | mmol/L |
| P | 1.29 | mmol/L |
| Fe* | 21 | µmol/L |
| Mn | 3.4 | µmol/L |
| Zn | 4.7 | µmol/L |
| B | 14 | µmol/L |
| Cu | 6.9 | µmol/L |
| Mo | 0.5 | µmol/L |
| EC | 1.4 | mS/cm |
| pH** | 6.1 |  |

\*Consists of 50% Fe-DTPA 3% and 50% Fe- EDDHSA 3% \*\*pH adjusted with Potassium hydroxide and Sulphuric acid

**Supplementary Table 4.**
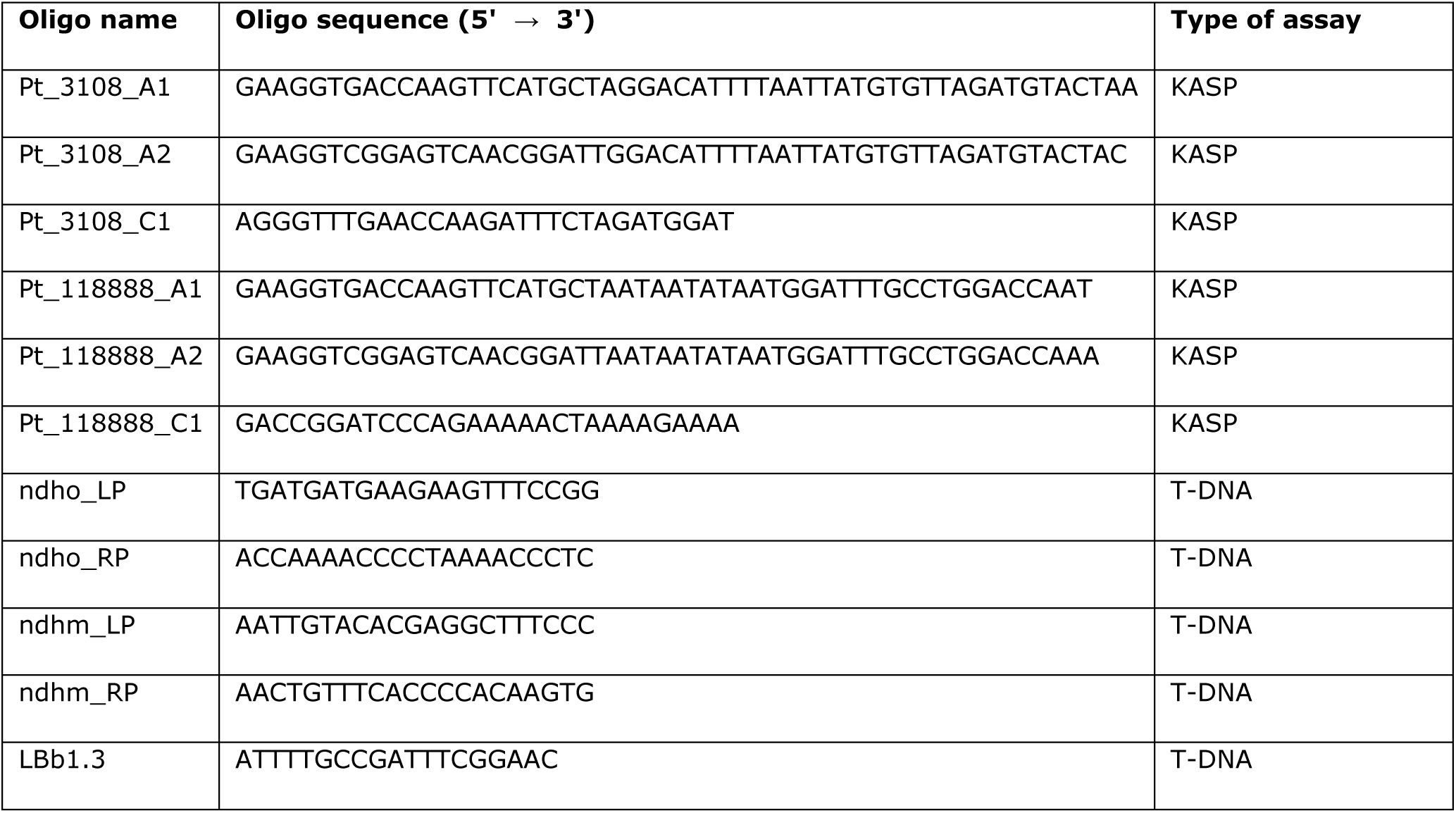
Primers used in this study. For the KASP primers the position on the reference genome TAIR10.1 is listed. For the TDNA primers, the left primer (LP), right primer (RP) and TDNA border primer (LB) is given.

## Supplementary Figures

**Supplementary Figure 1.**
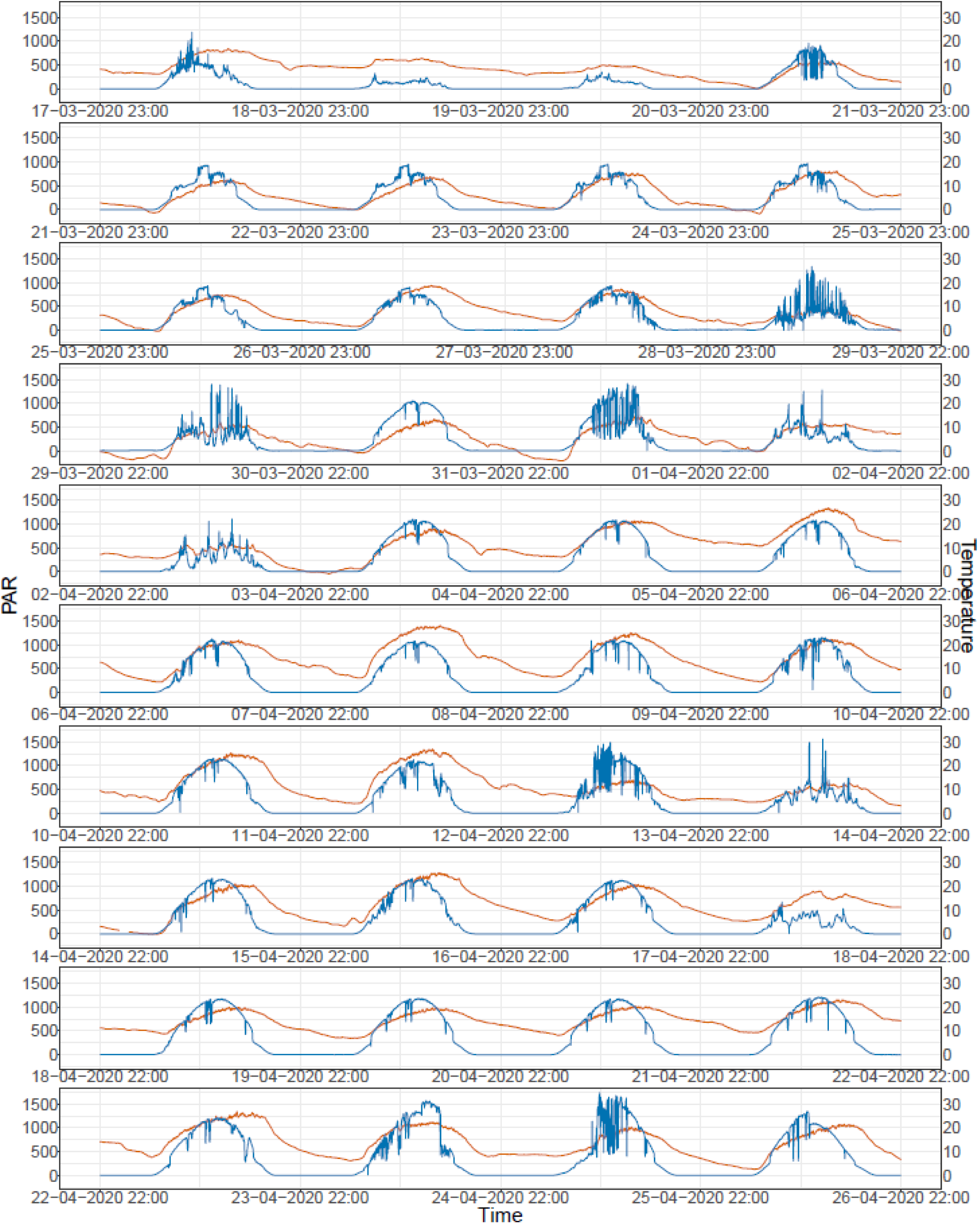
Light intensity (µmol m^-2^ s^-1^) and temperature for the semi-protected experiment in spring 2020.

**Supplementary Figure 2.**
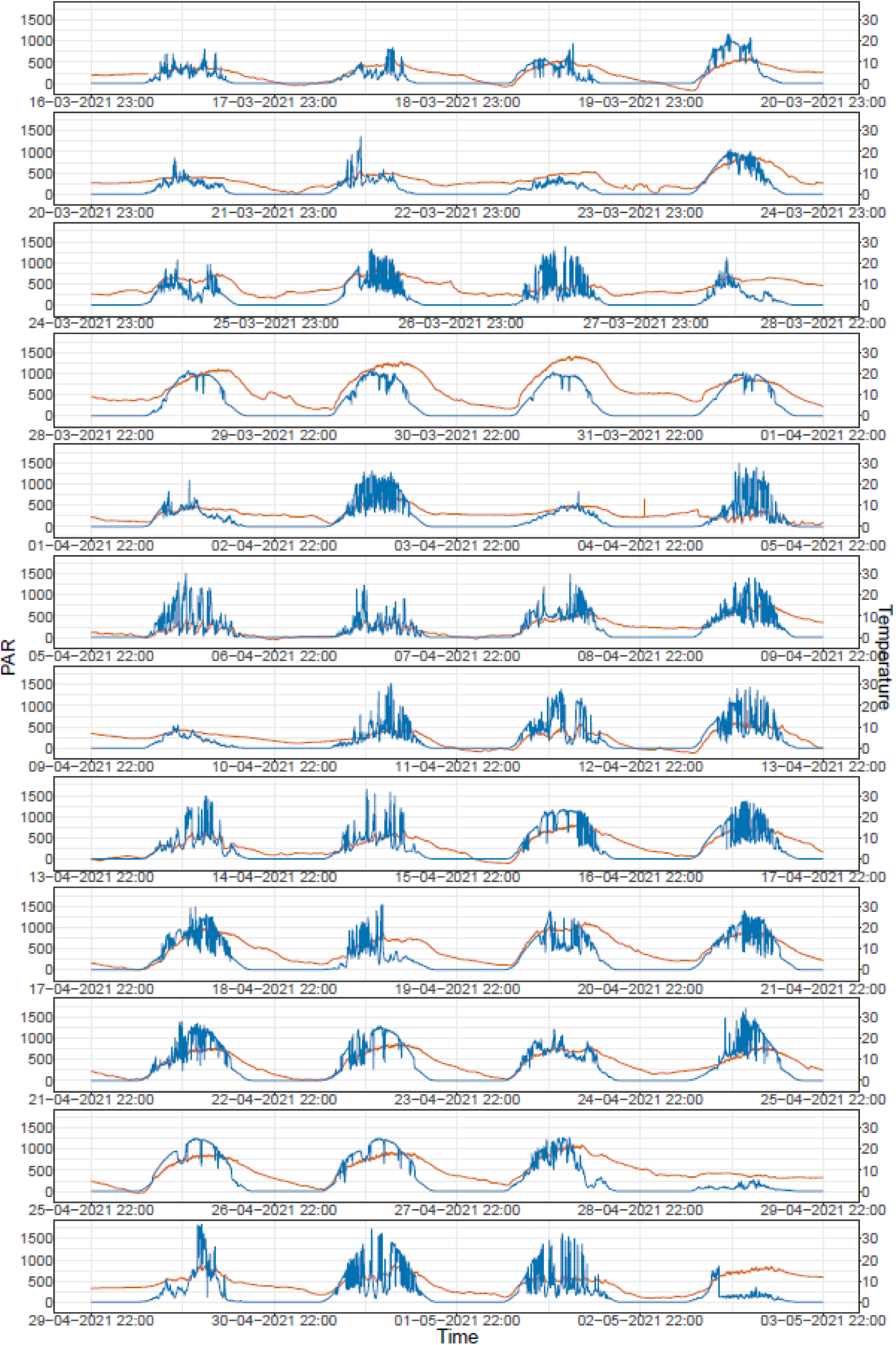
Light intensity (µmol m^-2^ s^-1^) and temperature for the semi-protected experiment in spring 2021

**Supplementary Figure 3.**
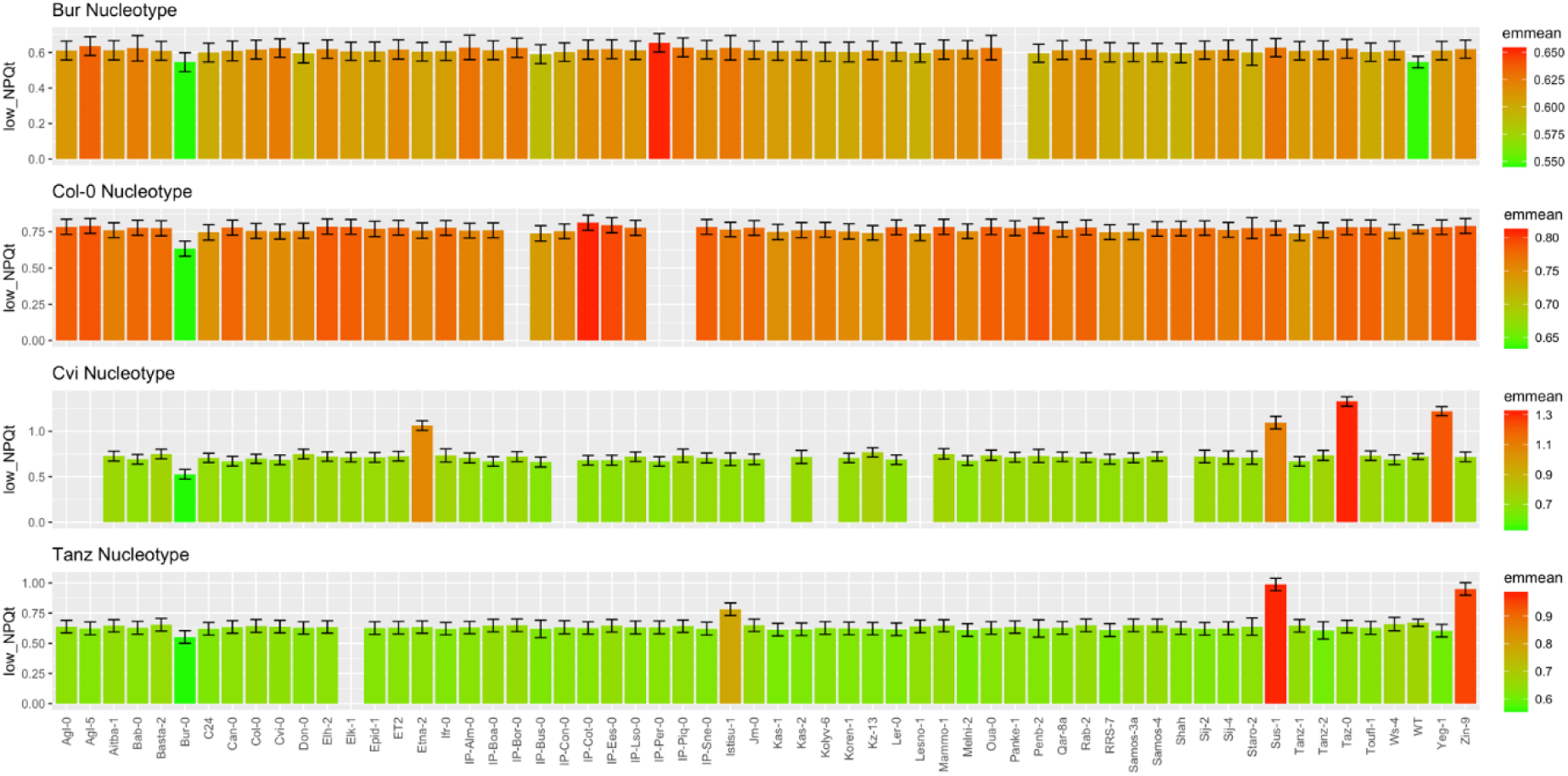
Combined analysis of spring 2020 and 2021 in semi-protected conditions for NPQt after 120s at 200 µmol m^-2^ s^-1^ light, after 1000 µmol m^-2^ s^-1^. The colours are scaled per nucleotype. The error bars represent the standard error of the mean.

**Supplementary Figure 4.**
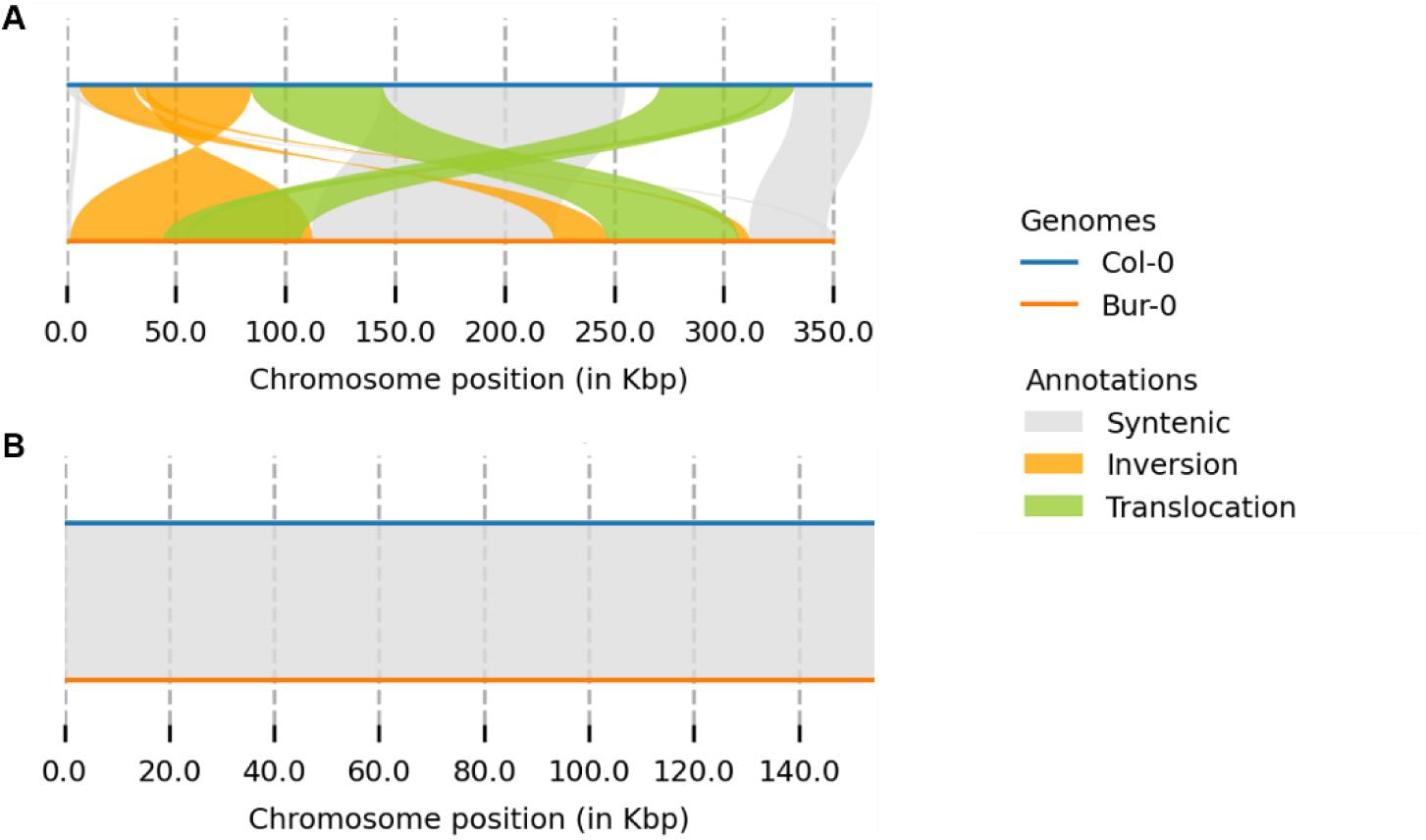
Structural differences between the de novo assembled organellar genomes of the Bur-0 and Col-0. A) The representation of the mitochondrial genome. B) The representation of the chloroplastic genome.

**Supplementary Figure 5.**
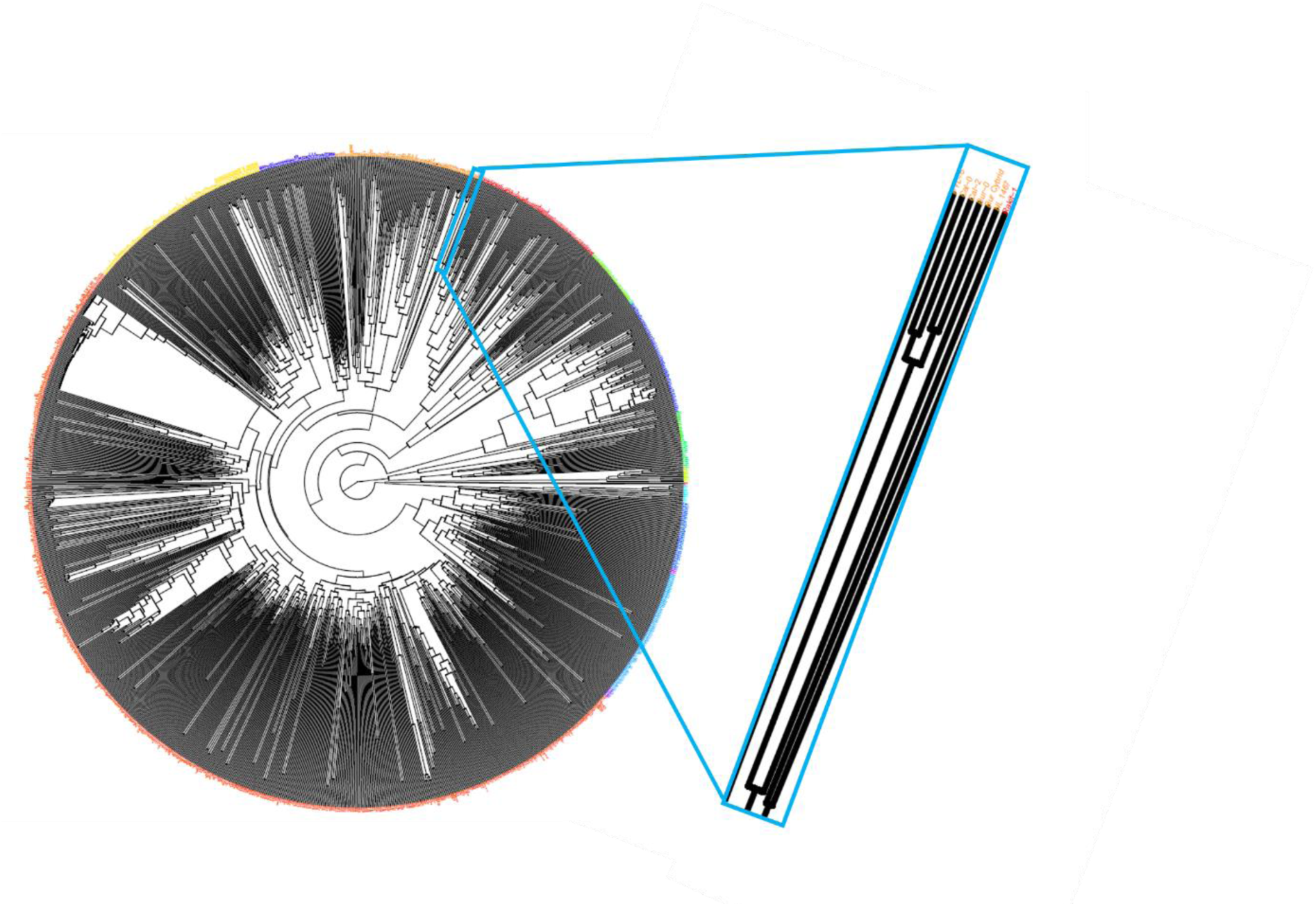
Hierarchical clustering of 1531 Arabidopsis thaliana accession (k=20), based on nuclear variation. Zoom in on section that shows accessions Bur-0, NL1467, Cal-0 and Cal-2.

**Supplementary Figure 6.**
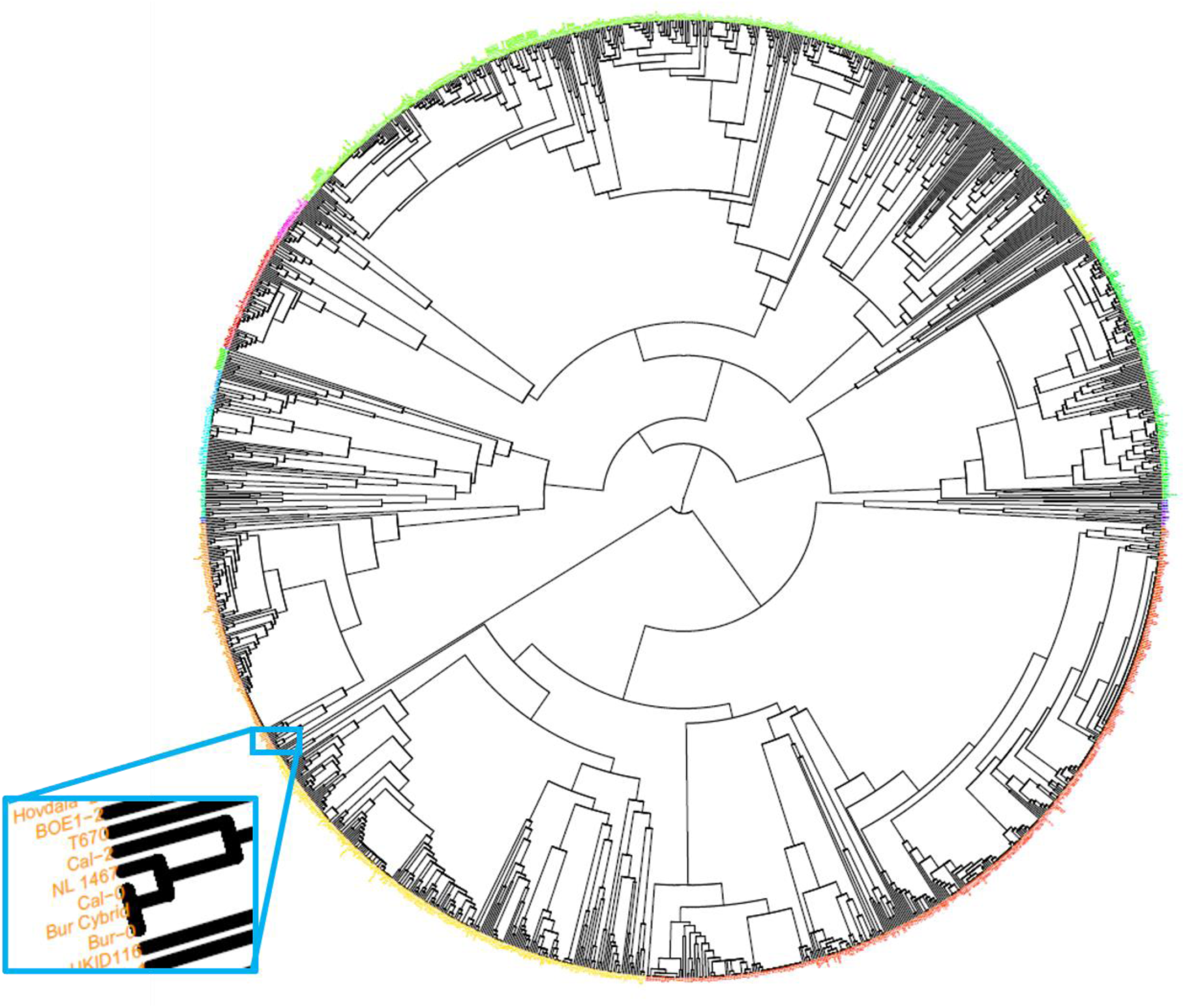
Hierarchical clustering of 1,531 Arabidopsis thaliana accessions (k=20), based on chloroplastic variation. Zoom in on section that shows accessions Bur-0, NL1467, Cal-0 and Cal-2. The rest of the orange cluster consists of the 117 accession which share the YCF1 and MATK SNP with Bur-0, NL147, Cal-0 and Cal-2.

**Supplementary Figure 7.**
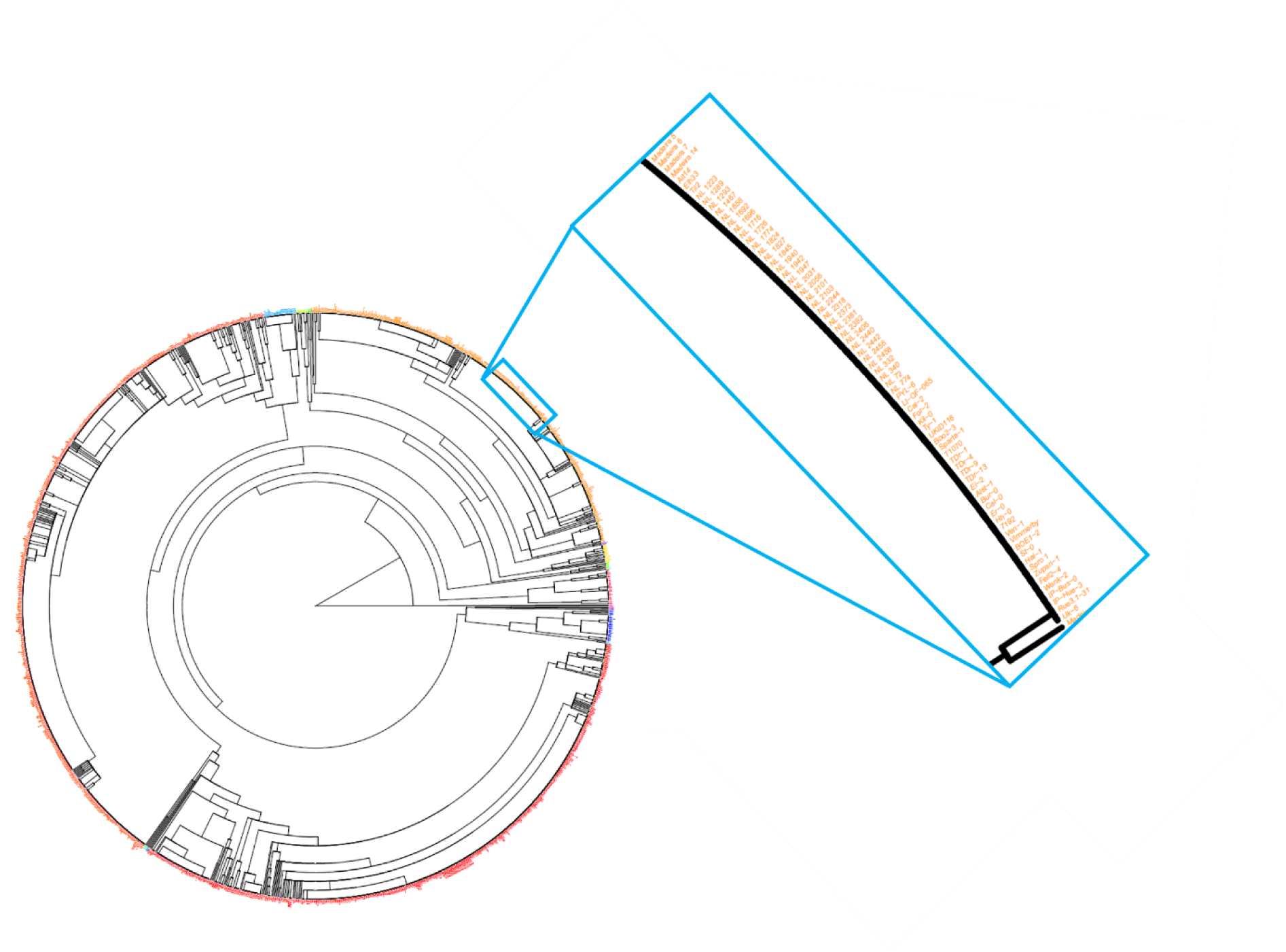
Hierarchical clustering of 1,531 Arabidopsis thaliana accessions (k=20), based on mitochondrial variation. Zoom in on section that shows accessions Bur-0, NL1467, Cal-0 and Cal-2.

**Supplementary Figure 8.**
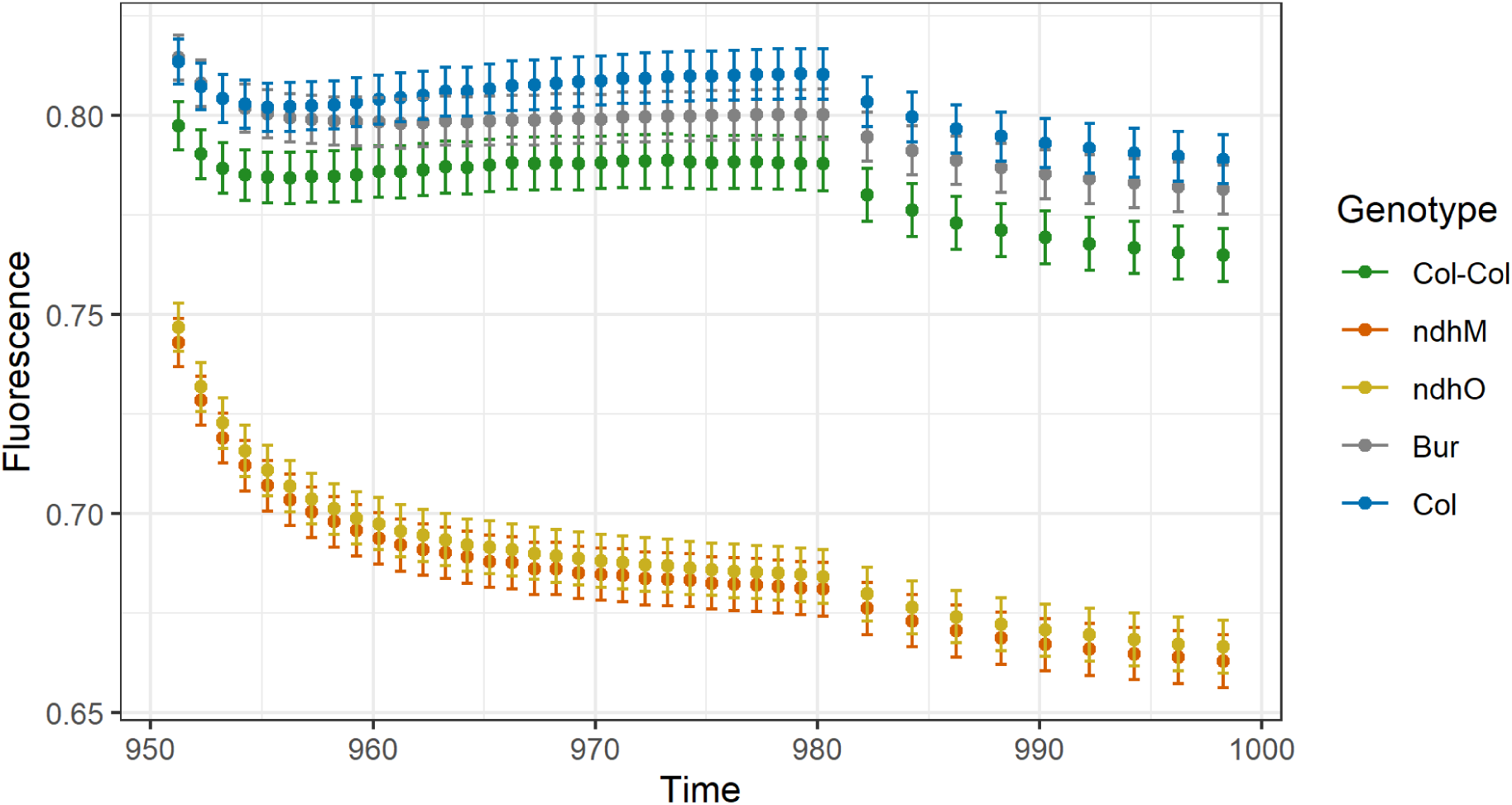
Post illumination fluorescence rise of after 200 µmol m^-2^ s^-1^, with t = 950 seconds the initiation of darkness. The Col-0 control is used (Col-Col) and the ndhm and ndho mutants. Also, the additive effect of the Bur-0 and Col-0 plasmotype is given, as averaged over the four nuclear genomes (Col-0, Bur-0, C24 and Ler-0).

**Supplementary Figure 9.**
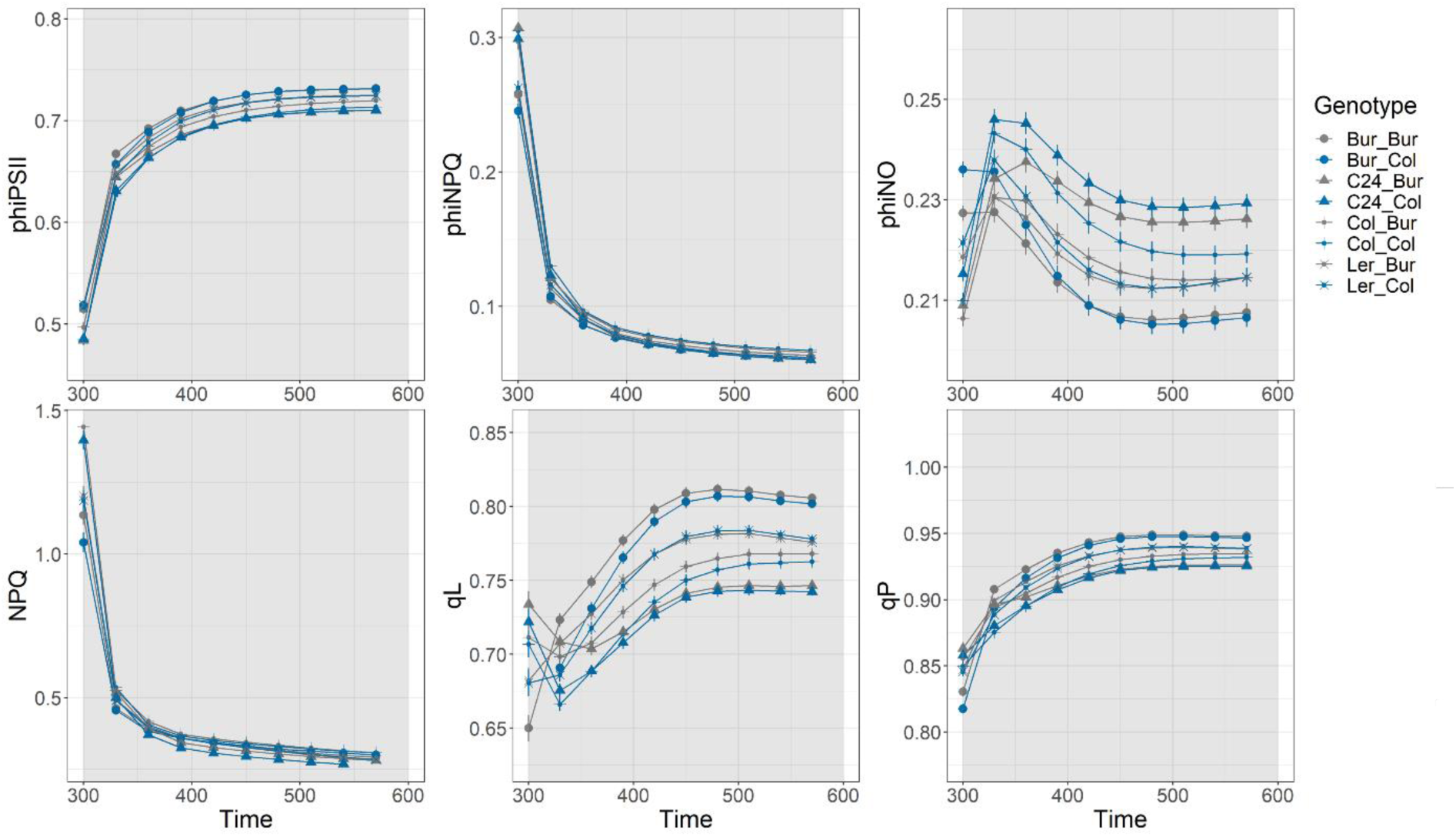
Phenotypic response of cybrids 50 µmol m^-2^ s^-1^, right after five minutes of 1,000 µmol m^-2^ s^-1^. At t=300 seconds the 50 µmol m^-2^ s^-1^ was initiated. The data as shown here is used as additive effects for the Bur-0 and Col-0 plasmotype and visualized in Figure 6.

**Supplementary Figure 10.**
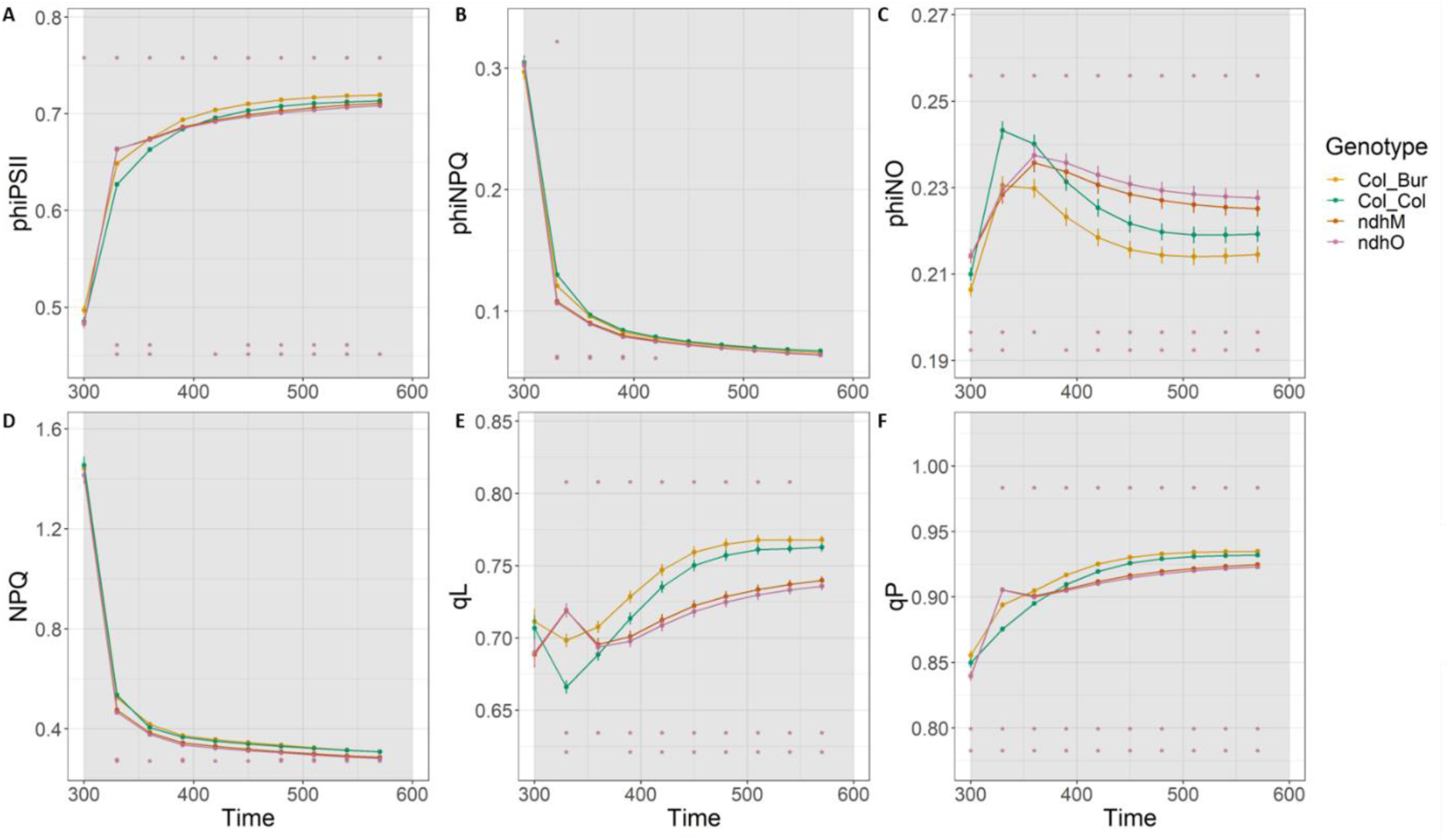
Phenotypic response of different genotypes 50 µmol m^-2^ s^-1^, right after five minutes of 1000 µmol m^-2^ s^-1^. At t=300 seconds the 50 µmol m^-2^ s^-1^ was initiated. The ndhm and ndho mutants are shown, and the control is Col-0 (Col_Col). To visualize the Bur plasmotype effect, without being influenced by a nucleotype effect, the cybrid with Col-0 as nucleotype and Bur-0 as plasmotype is given as well.

